# Single-vesicle tracking of α-synuclein oligomers reveals pore formation by a three-stage model modulated by charge, curvature, lipids and ligands

**DOI:** 10.1101/2025.04.10.648112

**Authors:** Bo Volf Brøchner, Xialin Zhang, Janni Nielsen, Jørgen Kjems, Daniel E. Otzen, Mette Galsgaard Malle

## Abstract

Neurodegenerative disorders like Parkinson’s disease (PD) pose significant health challenges. A major hallmark is the aggregation of α-synuclein into toxic oligomers (αSO) and fibrils. While many efforts focus on slowing disease progression, the molecular origins and mechanisms of αSO toxicity remain poorly understood, particularly its proposed link to membrane disruption. To address this, we have developed a single-vesicle analysis platform for direct and real-time measurements of αSO and membrane interaction. This allows us to show real-time translocation of dyes through αSO pores with single-particle resolution and single-channel electrical recordings for analyzing pore formation in planar lipid bilayers. Across methods, our data reveal evidence for a three-stage model for αSO and membrane interactions with initial membrane recruitment followed by partial pore insertion and subsequent full pore formation. Strikingly, while αSO recruitment was found to favor curved membranes, pore formation occurred more efficiently in less curved membranes, hence recruitment is divorced from a membrane charge-promoted reorientation and pore integration. Single αSO pore formations are undergoing multiple translocation steps hence pore formation is highly dynamic cycling back and forth from partial insertion to full pore formation. The dynamic nature of pore formation can be modulated by lipid charge, lipid headgroup class, and ligand binding. Our findings suggest that increased dynamic pore formation could implies increased membrane toxicity. Evidence for the three-stage model is important for future targeting strategies for blocking αSO mitigated PD-related cellular dysfunction and we envision the single-vesicle assay enables screening for ligands modulating the pore formation.

## Introduction

Membrane integrity is fundamental to cellular survival, serving both as a barrier and a dynamic platform for biological processes such as signaling, trafficking, and energy transduction. Membrane disruption can severely compromise cellular function, particularly in neurodegenerative diseases, where membrane damage has been implicated in the loss of neuronal function and cell death [1-3]. Related to this, aggregation of the protein α-synuclein (α-Syn) is central to both disease pathology and progression in the common neurodegenerative disorder Parkinson’s disease (PD) and other synucleinopathies. α-Syn is an intrinsically disordered protein hypothesized to be involved in synaptic vesicle regulation and trafficking [4]. However, misfolding and aggregation of α-Syn are associated with the formation of pathological protein inclusions called Lewy bodies [5] as well as soluble α-Syn oligomers (αSOs). These oligomers are considered a key factor in membrane disruption and subsequent neuron dysfunction, being more cytotoxic per mass than the more well-characterized fibrillar species [6]. While αSOs have been shown to compromise lipid membrane integrity, the mechanism of this event remains poorly understood [7-9].

The intrinsic heterogeneity of αSOs poses a significant challenge to elucidating the structure-function relationship. Unlike fibrils, which are highly uniform and give rise to atomically resolved structures, αSOs exist as a diverse population with varying sizes, conformations, and toxic potential [6, 10, 11]. Generally, αSOs can be subdivided into two distinct types: “on-pathway oligomers,” which act as intermediates in fibril formation, and “off-pathway oligomers” which arise through an alternative process that does not lead to fibril formation [6, 12-14]. *In vitro* studies have produced αSOs in both subdivisions with distinct structural and functional properties depending on the conditions under which they form. This variability on top of the intrinsic heterogeneity has hindered the development of a unifying model for how αSO-membrane interactions drive toxicity. This study focuses on the major toxic off-pathway oligomer, which forms spontaneously during fibrillation and accumulates as a stable and toxic species that does not lead to fibrils. Hence, they are believed to be both physiologically relevant and likely to be found in the brains of patients with PD [7, 15, 16]. Given the cytotoxicity of αSO, identifying high-affinity ligands for therapeutic and diagnostic applications is of great interest. One example of small ligands is nanobodies (NBs). NBs are modified variable domains from the heavy chain-only antibodies, which are another form of antibodies found in camelids [17, 18]. These have been suggested to have great therapeutic potential, i.e., due to their stability, high specificity, and hypothesized ability to cross the blood-brain barrier [19, 20]. We have recently produced two different NBs with high affinity and specificity for the major αSO species [21].

Previous studies have demonstrated that αSOs preferentially target negatively charged lipid membranes, such as those containing phosphatidylglycerol (PG) or phosphatidylserine (PS), over neutral lipid membranes like phosphatidylcholine (PC) [7-9, 22-26]. Negatively charged lipids are abundant in neuronal membranes, particularly in synaptic vesicles and mitochondrial membranes, making these systems highly relevant to PD pathology. Interestingly, while PS-containing membranes bind α-Syn efficiently, they exhibit less membrane leakage compared to PG-containing membranes [9, 23, 24]. This paradox underscores the complexity of lipid-specific interactions and suggests that membrane disruption is influenced not only by charge but also by curvature and headgroup structure.

αSO-driven membrane disruption is speculated to involve a partial insertion into lipid bilayers, which leads to their destabilization, allowing ions or other molecules to leak through [7, 9, 22-24, 27]. However, the details of this process, including how lipid composition and curvature modulate this disruption, remain unclear. Most studies rely on bulk techniques, such as the release of fluorescent dyes or ions from large unilamellar vesicles (LUVs) or small unilamellar vesicles (SUVs) [7-9, 22-25]. While these methods provide valuable insights, they average out the unavoidable heterogeneity from both the membrane vesicles and the inherently diverse αSO species, failing to capture the real-time dynamics and individual steps of membrane disruption [26, 28]. Single-vesicle techniques are essential for addressing these limitations. Total internal reflection fluorescence (TIRF) microscopy technique allows for real-time observation of protein-membrane interactions at the single-vesicle level [28, 29]. This bypasses ensemble averaging, revealing transient events and unraveling heterogeneous behavior invisible in bulk measurements.

Here, we report a single-liposome assay for direct and real-time measurement of αSO and membrane interaction. We have screened > 500,000 individual liposome-αSO interactions, providing the basis to build a biophysical understanding of the interactions in unprecedented detail. We propose a three-step model for αSO and membrane interactions with initial membrane recruitment followed by partial insertion and subsequent full pore formation. Recruitment preferentially occurs on highly curved membranes, while pore formation is favored on lower-curvature membranes with less tension. Pore formation is observed as highly dynamic, cycling back and forth from partial insertion to full pore formation, which is modulated by charge, lipids, and αSO-binding nanobodies. Our findings set the stage for future mechanistic investigations of αSO-membrane interactions and provide a platform for screening oligomer-membrane and ligand interactions, overcoming inherent heterogeneity. Understanding the modulable and dynamic pore formation might pave the way to mitigate αSO-induced membrane dysfunction.

### Direct observation of charge-promoted recruitment of individual αSOs to single-liposome vesicles

To address how individual αSOs interact with lipid membranes, we established a single-vesicle membrane assay and exposed it to the major cytotoxic off-pathway αSO species [Fig. 1a] that forms spontaneously during aggregation-inducing conditions and is known to form a pore-like structure [7, 13, 30, 31]. The αSOs were isolated from the monomer species by size exclusion chromatography [32] as a large (∼450 kDa), appearing as a clear band on a SDS-PAGE gel [Fig. 1b and Supplementary Fig.1].

**Figure 1.**
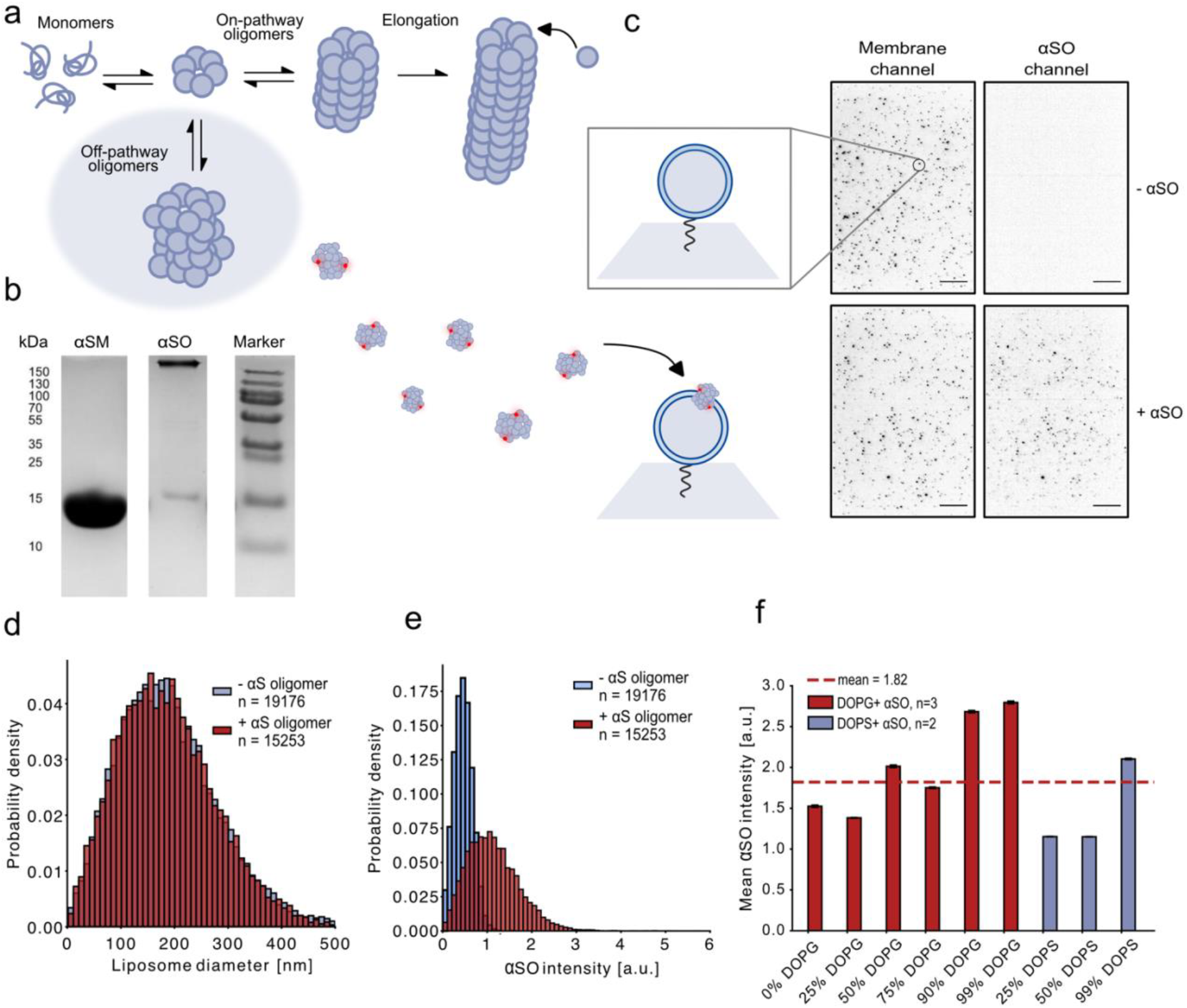
Single liposome detection shows charge-dependent recruitment of a cytotoxic off-pathway αSO. (a) Schematic representation of α-Syn aggregation. Derived from an initial aggregation model event, cytotoxic off-pathway αSO can be formed that do not continue their fibrillation to form fibrils. (b) SDS-PAGE of purified monomeric α-Syn and off-pathway toxic αSO. (c) Single-particle assay for αSO recruitment. Biotinylated liposomes are immobilized on a passivated glass surface where an incorporated lipidated-fluorescent dye in the liposome membrane allows detection of it in the membrane channel. In the bottom panels fluorescently labeled αSO is added and can be detected in the αSO channel. Colocalization in the two channels allows for quantification of the amount of αSO bound to each single liposome. (d) Histogram showing no change in liposome size distribution before and after the recruitment of αSO. (e) Histogram showing a clear intensity shift in αSO intensity before (blue background signal) and after the membrane recruitment of αSO (red). (f) αSO associations to more than half a million single liposome membranes in response to increasing charge (increasing DOPG content) and different lipid composition (DOPS and DOPG lipids). Data is recorded in n=3 biological replicates for DOPG vesicles and n=2 for DOPS vesicles, error bars show the error propagated SEM.

Next, we produced SUVs (liposomes) containing 0.5 % DSPE-PEG(2000)-Biotin, allowing us to tether them to a PLL-g-PEG-biotin-covered glass surface via a neutravidin linker. The liposomes also contained 0.5% ATTO-DOPE, allowing imaging with a high signal-to-noise ratio with a total internal reflection microscope (TIRF). Forty-eight micrographs were taken before and after the addition of ATTO655-labeled αSO [Fig. 1c]. A total of 578863 liposomes were analyzed across the 3 replicates for each liposome condition. Fig. 1c shows a representative field of view of both the liposome channel and the αSO channel before and after the addition of the αSOs, showcasing a clear membrane association of the αSOs. All liposomes were signal-integrated to extract both nanometer-precise localization and membrane intensity using an in-house Python script. The single vesicle membrane intensity units were converted to liposome diameters from the mean integrated intensity normalized to the mean vesicle size from nanoparticle tracking analysis (NTA) of the suspended liposome for each liposome preparation [28, 33] [Supplementary Fig. 2]. Measurements of the size distribution of liposomes showed no change upon addition of αSO [Fig. 1d and Supplementary Fig. 3-4]. The vesicle localization script allowed us to co-localize all membrane-associated αSO signals, enabling quantification of αSO on each liposome.

Here, we integrated the αSO signal and compared the intensity distributions before and after the addition of labeled αSO. Using a Kolmogorov–Smirnov test (KS-test), we found a significant shift in the signal distributions after αSO recruitment [Fig 1e and Supplementary Fig. 5-6]. When no liposomes were tethered to the glass slide, we saw no αSO bound to the surface, demonstrating that binding is membrane dependent, and our assay has minimally false positive association or false membrane signal integration [Supplementary Fig. 7].

To test the charge-dependent membrane recruitment of αSO, we prepared a series of increasingly anionic liposomes, containing 0-99% DOPG and 0-99% DOPS. A clear tendency that recruitment increases with negative charge was observed [Fig. 1f]. However, neutrally charged membranes (0 % DOPG) still distinctly recruit αSO. DOPS liposomes show lower recruitment than DOPG liposomes, indicating an important role for the headgroup. Also, DOPS membranes show more cooperative (all-or-nothing) behavior, with highly significant elevated recruitment at 99% DOPS. This is consistent with calcein-release assays, where more negatively charged membranes lead to a higher release of dyes and DOPG/POPG-containing vesicles are more destabilized by αSO than DOPS/POPS-containing vesicles [7-9, 22-25].

### Membrane charge- and curvature-promoted recruitment of αSO

A mechanistic understanding of αSO recruitment requires deconvolution of lipid composition, charge, and curvature, which is averaged out due to heterogeneity in bulk studies. Our single-vesicle analysis addresses this limitation by resolving the relationship between individual liposome sizes [Fig. 2a] and αSO recruitment, enabling us to screen membrane properties and transform molecular stochasticity into an experimental advantage. Here, surface-tethered liposomes were integrated in the membrane intensity channel and colocalized with the membrane-bound αSO signal, enabling the calculation of αSO density for each liposome and its curvature [Fig. 2a], given the inverse relationship between curvature and vesicle diameter[34, 35]. This allowed us to screen single curvature dependent recruitment quantified across three biological replicates yielding a total of 362787 liposomes [Fig. 2b, c and supplementary Fig. 8,9]. Here the αSO density (*D*_*αSO*_) was plotted against liposome size and binned on 10 nm intervals ranging from 25nm to 400 nm in diameter and fitted with an offset power function: *D*_*αSO*_ = *D*_0_ + *β* ∗ (*Dia*_*liposome*_) ^*α*^, for each membrane condition. The exponent α is the rate of curvature dependency, D_0_ is the density offset while the scaling factor β quantifies the αSO’s sensitivity to curvature dependency [33].

**Figure 2.**
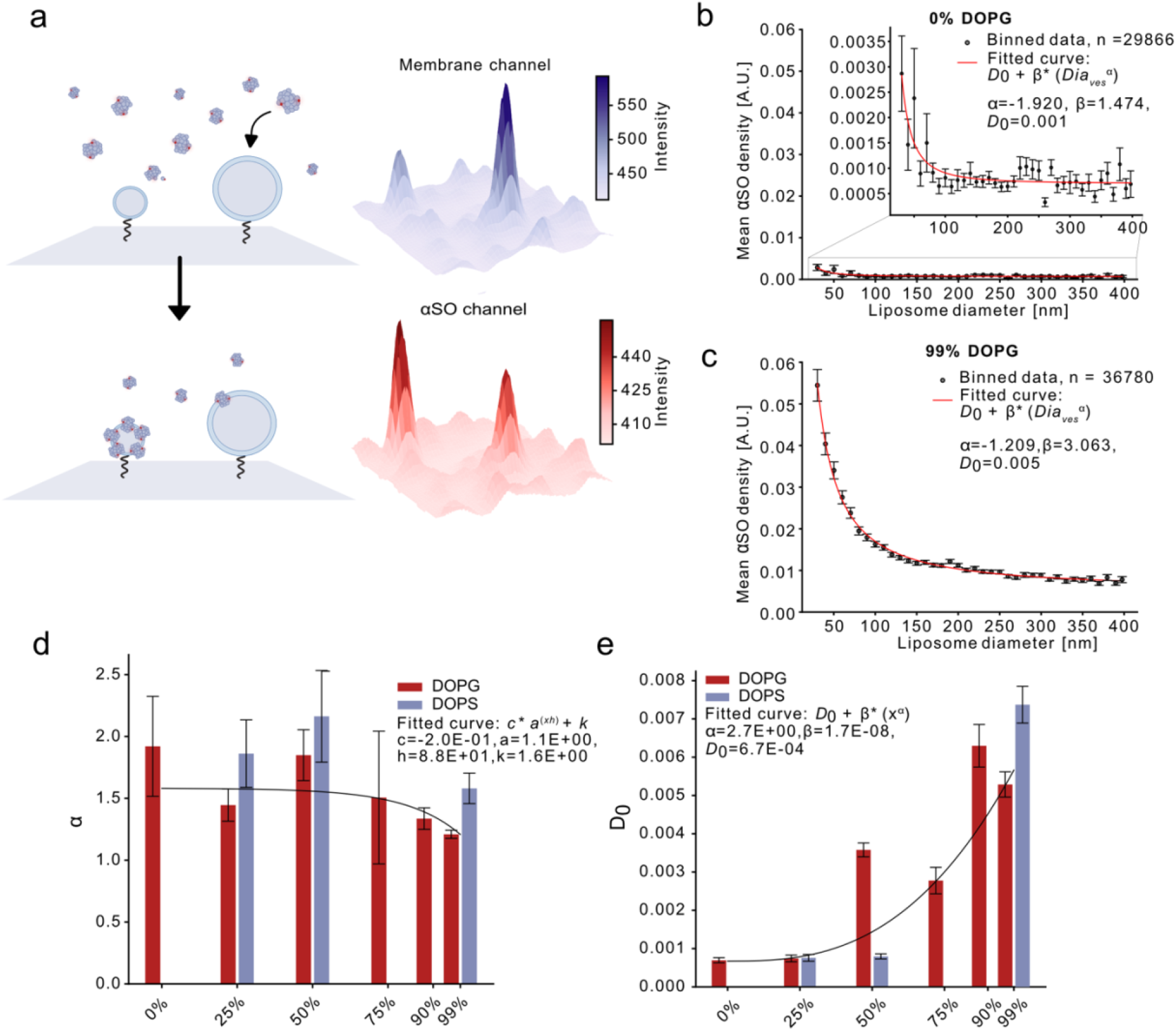
Single vesicle detection shows higher αSO density with increasing curvature. (a) The co-localization of immobilized liposomes and associated αSO enable direct measurement of the amount or density of αSO per liposome. The Inevitable heterogenous liposome size distribution allows for screening of membrane-dependent behavior such as curvature. Representative images in c and d show larger liposomes, hence less curved, display lower αSO density. (b) Liposome diameter as a function of membrane density of αSO for liposomes with 0% DOPG. The relation is fitted to a recruitment power function (D = D_0_·β·D_vesicle_^a^) showing a decrease in recruited density for increasing liposome size. The insert shows the steep growth rate recruitment for small liposomes. (c) Liposome diameter plotted against membrane density of αSO for liposomes with 99% DOPG. The relation is fitted to a power function which shows a decrease in recruited density for increasing liposome size. However, this shows a less strong power function dependency than for the 0% DOPG liposomes. The data in (b,c) is binned per liposome size for each 10nm with error bars showing the SEM. (d) Liposomes with increasing charges for both DOPG and DOPS vesicles are all measured at a single-vesicle level and fitted to the recruitment’s power function. The exponent of the fit α, which refers to the strength of the recruited density for curvatures, depicts a negative correlation for increasing anionic membrane charge. Hence, lower-charged vesicles are more curvature-dependent in the recruitment of αSO. (e) The density offset from the recruitment’s power function for increasing charges for both DOPG and DOPS vesicles shows an increasing offset upon increasing membrane charge. The data suggest that the higher charged liposomes overall recruit more αSO, but the recruitment is less dependent on curvature. Error bars in (d,e) show std.

Neutrally charged liposomes showed a strong curvature promoted recruitment with a rate of curvature dependency factor α = -1.920 ± 0.40 [Fig. 2b]. The density offset value, D = 0.001 ± 6.69 ∗ 10^−5^, is reflecting less binding on flat membranes and overall low αSO density. The high α and low D_0_ values reflect low αSO recruitment on less curved, neutral membranes, consistent with the reported lack of binding to GUVs [24]. Negatively charged membranes (99% DOPG) showed a significantly higher density offset value, D_0_ = 0.005 ± 0.0003 [Fig. 2c] reflecting a significantly stronger αSO recruitment. Surprisingly, the curvature dependency (α) is significantly lower than neutrally charged membranes with α-= -1.209 ± 0.033. Overall αSO recruitment depends on D_0_, α, and β, quantified by integrating the αSO density across all liposomes. For 99% DOPG liposomes, the integral is 5.296, whereas neutrally charged liposomes exhibit a substantially lower value of 0.451. To fully investigate the charge-dependent single-vesicle recruitment of αSO, we prepared a series of increasingly anionic liposomes, containing 0-99% DOPG and 0-99% DOPS [Supplementary Fig. 8 and 9] and extracted α and D_0_ values to compare the curvature promoted recruitment across the membrane compositions [Fig. 2d and 2e]. Here the curvature dependency (α) for DOPG-containing liposomes is found to follow an exponentially decreasing function with increasing anionic charge, demonstrating that neutrally charged membranes are more curvature-dependent and the dependency decreases exponentially with increasing negative charges. DOPS liposomes showed stronger curvature-promoted recruitment, which is particularly evident at a high negative charge, with 99% DOPG exhibiting an α-value of -1.209 ± 0.033, compared to the significantly higher α-value of 1.580 ± 0.123 for 99% DOPS. Despite this disparity, DOPS vesicles show a similar overall trend, with curvature dependency decreasing as membrane charge increases. The offset density [Fig. 2e] increases with membrane charge for both lipid compositions, with a power-law correlation, showing steep charge-dependent recruitment. DOPS liposomes exhibit a steeper increase in offset, indicating a stronger and more cooperative charge dependency compared to DOPG liposomes.

To our knowledge, this is the first quantitative description of αSO recruitment using single-vesicle measurements, avoiding ensemble averaging and enabling direct comparison of lipid headgroup, charge, and curvature effects. Our data demonstrate that neutral membranes can recruit αSO, but recruitment is highly curvature-promoted, likely due to lipid packing defects exposing hydrophobic tails. Recruitment increases significantly with higher anionic membrane content, however, we find that also the lipid headgroup play an important role, as αSO recruitment to DOPS membranes is more sensitive to charge and curvature than to DOPG membranes.

### Single-vesicle fluorescent and single-channel electrical recordings show αSO pore formation and small-molecule translocation

To gain deeper insight into the interaction between αSO and lipid membranes, we employed a single-vesicle setup to directly measure how αSOs permeabilize or perforate membranes. While previous studies have demonstrated that αSOs induce leakage or disruption, particularly in anionic vesicles, these findings are based on bulk measurements that cannot disentangle specific interaction mechanisms—such as lysis or pore formation—or elucidate the precise nature or kinetics of the insertions [7, 24]. To address this limitation, we encapsulated ATTO-655 carboxy in 0.5% ATTO-488-DOPE membrane labeled liposomes, enabling synchronous recording of both vesicle membrane and lumen dye [Fig. 3a]. Liposomes were immobilized and recorded for 5 hours with a temporal resolution of 1 frame per minute. After 7 minutes, 0.31 µM αSO was added using an automated peristaltic pump setup. In-house automated software was used to track and extract spatiotemporal information for individual liposome monitoring each membrane and their lumen molecules. Notably, as seen in intensity-time trajectories [Fig. 3b] the membrane signal (blue) remained stable throughout the 5-hour observation period, while the lumen dyes showed two distinct step-like events, occurring at 15 and 75 minutes for this particular particle, indicating full pore insertion followed by partial insertion and back to full insertion leading to two subsequent translocation steps of the dyes. This indicates that lumen dye release is mediated by αSOs insertion and pore formation into the membrane as liposome membranes retain the structural integrity [Supplementary Fig 10 for additional trajectories].

**Figure 3.**
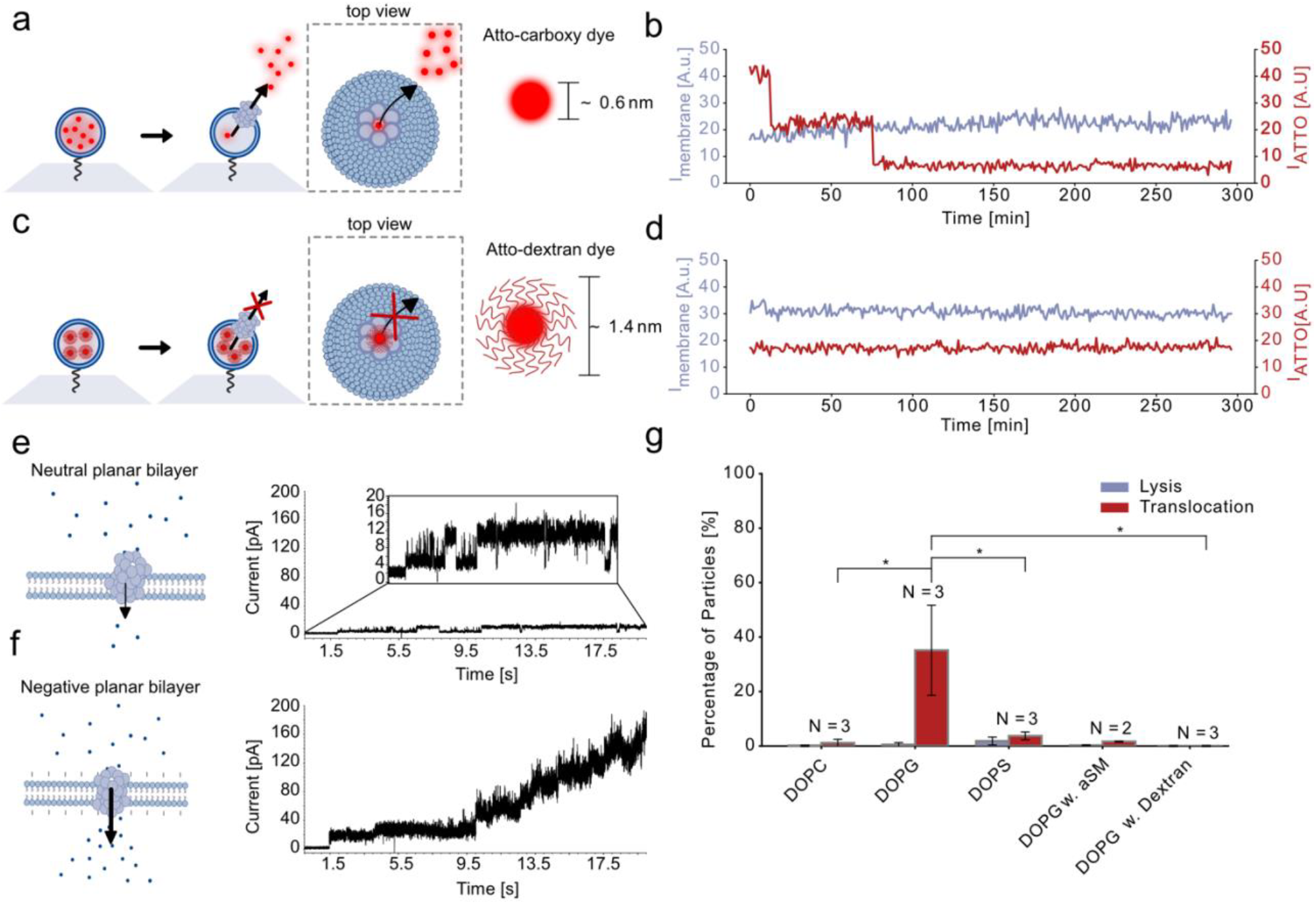
Real-time single-vesicle fluorescent and single-channel electrical recordings demonstrating αSO pore formation and small-molecule translocation. (a) Liposomes are membranes labeled with ATTO488-DOPE and ATTO655 carboxy are encapsulated, allowing simulations and real-time recording of both the membrane and the lumen dyes using a TIRF microscope. αSOs were injected into the microscope chamber after 10 frames of recording. (b) Representative time trajectories were obtained from real-time tracking and integration of both the membrane (blue) and lumen dye (red) intensities. Here, the trajectory shows a stable liposome membrane and a dynamic pore formation with two distinct open pore formations, which allows for the translocation of dyes. (c) Liposomes are membrane labeled with ATTO655-DOPE and the larger ATTO 488-dextran 4kDa dye was encapsulated. (d) Representative time trajectories displaying a stable membrane and no translocation of ATTO 488-dextran. (e) Representative current trace showing the partial insertion of αSO into neutral planar lipid bilayers. Prior to recording, 1.4 ug of αSO was added to each side of the chamber. The neutral-charged planar lipid bilayer was formed using DphPC. (f) Representative current trace showing the full insertion of αSO into negative-charged planar lipid bilayers. Prior to recording, 1.4 ug of αSO was added to each side of the chamber. The negative-charged planar lipid bilayer was formed using DphPG. (g) Statistics from thousands of single liposome tracking, as explained in (b,d). Interestingly the activity of αSO in charged 99% DOPG vesicles shows a significant elevated pore formation. The monomer shows no pore activity, which is confirmed by electrical recordings. Finally, the translocation of dextran is not present, suggesting the translocation is through the small pore-like structure of the αSO. Asterisks (*) in (g) indicate a p-value <0.05 and reflect a significant difference based on a Mann-Whitney U test.

To confirm that only small molecules can translocate through αSO pores, we loaded liposomes with atto488-dextran, a dye with a hydrodynamic diameter of approximately 1.4 nm [Fig. 3c]. A representative trajectory in Fig. 3d shows stable intensity in both liposome membrane (blue) and encapsulated cargos (red) over a 5-hour observation period, indicating no translocation of the larger molecule [Supplementary Fig 11 for additional trajectories]. This suggests that larger molecules exceed the dimension of αSO pores, consistent with pore formation of a defined size rather than random permeabilization. Additionally, we measured small molecule ATTO-655 carboxy translocation in the presence of αS monomers. Here liposome remained stable, and no dye translocation was observed [Supplementary Fig 12], like the control with buffer alone [Supplementary Fig 13]. These results demonstrate that molecular translocation indeed occurs through the αSO pore.

To further confirm αSO pore formation, we employed a technique for measuring ionic conductivity using single-channel electrical recording setup on planar lipid bilayers [36, 37]. On a neutral planar lipid bilayer (using DphPC), αSO perforation led to ionic flux with an open pore current of 7.67 ± 2.08 pA at an applied voltage of 20 mV, which indicated the insertion of small pores. A characteristic current-time trajectory representing αSO insertion into a neutral lipid bilayer is shown in Fig. 3e [Supplementary Fig. 14 for additional trajectories]. The amplified current trace showed a single-step current increase, indicating limited pore insertion activity of αSO in a neutral lipid membrane. In contrast, the same amount of αSO (1.4 µg) induced a stepwise current increase and caused large ionic conductivity for negatively charged bilayers (using DphPG) [Fig 3f and Supplementary Fig. 15]. This observation suggests that αSO more efficiently inserts itself into negatively charged lipid membranes. Interestingly, the presence of CaCl_2_ did not significantly influence αSO pore insertion activity in either neutral or negatively charged planar lipid bilayer [Supplementary Fig. 16 and 17]. Furthermore, control experiments using the same amount of αS monomers (1.4 µg) on the negatively charged bilayer showed no insertion activity, confirming that the observed increase in ionic conductivity was specifically due to αSO pore formation [Supplementary Fig. 18].

To resolve the role of differently charged lipids in αSO pore insertion, we prepared 3 different types of liposomes using one neutrally charged (DOPC) and two anionic (DOPG and DOPS) lipids, encapsulated ATTO655-carboxy, and recorded in real time yielding thousands of single-vesicles (Supplementary Fig 19). In neutrally charged membranes, only a small fraction of vesicles exhibited translocation (1.14 ± 1.34%), and 0.18 ± 0.14% underwent lysis [Fig. 3g]. In contrast, vesicles with anionic DOPG membranes showed a dramatic increase in pore formation, with 35.16 ± 16.55% of vesicles exhibiting translocation, while only 0.52 ± 0.73% underwent lysis. Interestingly, anionic DOPS vesicles exhibited significantly lower pore insertion (3.71 ± 1.43% translocation) while a substantial fraction of these vesicles (1.83 ± 1.46%) underwent lysis, showing a higher propensity for membrane disruption compared to DOPG vesicles. Real-time recordings of αSM revealed minimal activity, with 1.67 ± 0.04% of vesicles showing translocation and 0.24 ± 0.03% undergoing lysis. As a control, vesicles containing large dextran-coupled dye encapsulated in negatively charged DOPG membranes exhibited no detectable translocation (0.0 ± 0.0%), confirming that dye release occurs through αSO pores. Collectively, our findings reveal that αSO can bind to both neutrally and negatively charged membranes, however, only the negative membrane charges activate pore formation. Additionally, we find subsequent step-like translocation of lumen dyes for single-vesicles. This suggests a novel three-stage interaction model involving initial αSO recruitment to the membrane surface then cycles between partial insertion followed by full pore formation. While the mechanism of membrane permeabilization by αSOs remains debated [8, 38], our results, in agreement with previous studies, demonstrate that binding alone does not necessarily lead to membrane rupture [24]. Our data suggests that pore formation in SUVs predominates over membrane lysis.

### The dynamics of αSO pore formation are sensitive to lipid charge and headgroup type

The time-resolved recording of dye translocation at single-vesicle level allowed us to investigate the effect of lipid and curvature for αSO pore formation. Here we find that pore formation is highly dynamic at the individual oligomer level cycling reversibly back and forth between a partial insertion stage and a full pore forming stage. We find that the dynamic or frequency of pore formation is sensitive to both lipid charge and headgroup type.

Again, we investigated neutral membranes (DOPC) and two anionic membranes (DOPG and DOPS) yielding thousands of vesicles and classified each vesicle as either i) non-reacting, ii) translocating (pore formation) or iii) lysed (simultaneous lumen and membrane disruption). The classified event was plotted as a function of the liposome diameter [Fig. 4a]. Neutrally charged liposomes (red) showed no curvature preference for pore formation, however it should be noted that the number of translocations is small as negative charges are found crucial for active pore formation. Notably, for negatively charged DOPG liposomes (blue), pore-forming events are highly significantly shifted toward larger, lower-curvature liposomes (P-value = 5.6·10^−30^ based on a one-sided Mann-Whitney U (MWU) test and P-value = 4.4·10^−16^ based on a Kolmogorov Smirnov (KS) test). A subset of the liposome population also underwent lysis, predominantly among larger liposomes (P-value = 0.01 MWU-test and P-value = 0.048 KS-test). In contrast, DOPS liposomes (green) show a smaller fraction of pore-forming events, however still predominantly in significantly larger liposomes with lower curvature (P-value = 1.2·10^−08^ MWU-test and P-value = 2.4·10^−08^ KS-test). Also, lysis is significantly associated with larger liposomes (P-value = 4.0·10^−16^ MWU-test and P-value = 2.4·10^−08^ KS-test).

**Figure 4.**
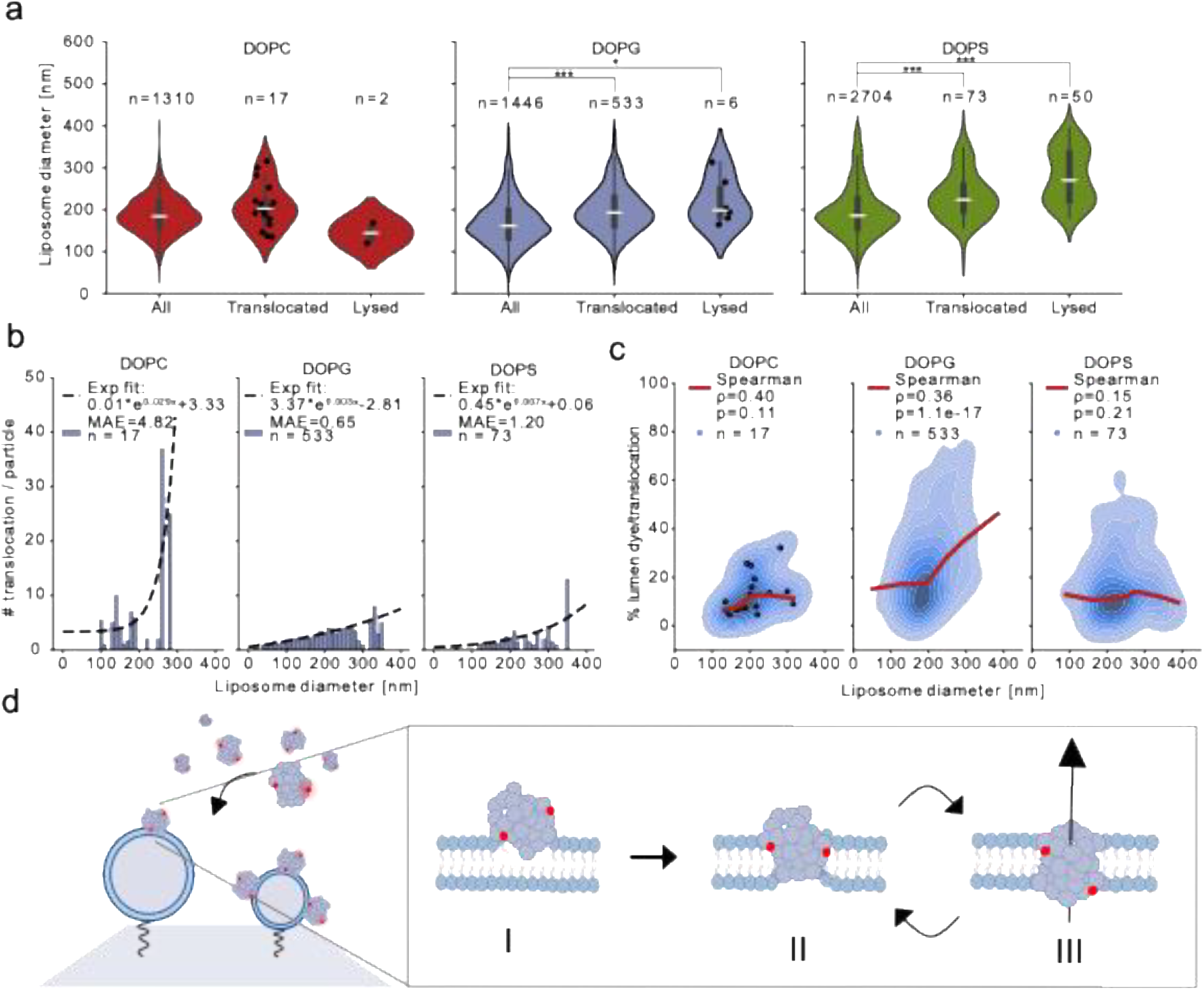
Single-vesicle recording shows αSO pore activity is regulated by the membrane lipids. (a) Distributions of αSO pore activity for neutral DOPC and charged DOPG and DOPS liposomes plotted against the liposome diameter. For the neutral DOPC liposomes (red), no curvature or size preference was found and only 0.65% and 0.08% of the liposomes were observed to have αSO pore translocations of dyes or lysed in the presence of αSO, respectively. For negatively charged DOPG liposomes (blue), pore-forming events were highly significantly shifted toward larger liposomes with lower curvature. A subset of the liposome population underwent lysis, also predominantly among larger liposomes. In contrast, DOPS (green) liposomes exhibited a smaller fraction of pore-forming events, though this fraction is likewise significantly enriched in larger liposomes. Lysis was more prevalent in DOPS liposomes and is similarly associated with larger liposomes. Asterisks indicate the following p-values with significance * = 0.05, ** = 0.01, *** = 0.005 based on a one-sided Mann-Whitney U test. (b) Quantification of the number of step-like translocations of dyes through the αSO pores. Here DOPG vesicles showed a linear distribution of the number of leaks per liposome. In contrast, DOPS and DOPC vesicles showed an exponentially increasing number regarding vesicle diameter. (c) The quantity of lumen dyes translocated in a single translocation event. DOPG vesicles show a positive correlation of % translocated with liposome diameter. For DOPS and DOPC liposomes we find no correlation. (d) Our results suggest a new three-stage model for αSO and membrane interactions with initial membrane recruitment preferably in smaller more curved vesicles followed by a reversible pore formation preferably in larger less curved membranes which strictly depends on membrane charge. Hence αSO recruitment to the membrane is decoupled from charge-dependent pore formation.

Next, the time-resolved single-vesicle level provided insight into an important aspect of αSO-membrane interactions, namely the dynamics in pore formation in the form of the frequency of pore formation or translocation turnovers as a function of liposome size [Fig. 4b]. The observed time-resolve on/off pore formation suggests that the oligomer is firstly recruited to the membrane followed by cycling back and forth between partial insertion and subsequent full pore formation, which we can observe as on/off translocations through the pore. We observed that individual liposomes can undergo up to 37 cycling events between being partially and fully inserted, where larger liposomes show a higher number of translocations. DOPG vesicles show a linear distribution of the number of translocations with increasing liposome sizes, whereas DOPS and DOPC vesicles show a slightly and more exponential relationship, respectively. Furthermore, for each translocation event, we quantified the quantity of the lumen [Fig. 4c]. Here DOPG vesicles show a positive correlation of % translocated with liposome diameter, hence larger liposomes are translocating a larger portion of dyes in a single translocation (p-value = 1.1·10^−17^ based on spearman correlation), suggesting the presence of larger or more stable pores for flat membrane. Conversely, DOPC and DOPS vesicles show no correlation in translocation as a function of liposome sizes. This might be due to less rotational or penetration freedom for the αSO due to less preferred membranes and charge. These findings suggest that pore formation is highly dynamic, with larger, less-curved liposomes exhibiting increased dynamic favorability. Moreover, pore formation is strongly lipid-dependent, being more pronounced in DOPG vesicles, where pores also appear to be more stable compared to those in DOPS vesicles. These findings might be of great biological relevance, as PG lipids are found in mitochondrial membranes, and PS lipids are found in synaptic vesicle membranes [39]. Overall, our findings reveal that while αSO recruitment is curvature-promoted, pore formation dominates significantly in lower-curvature membranes, hence the two steps are decoupled, supporting our proposed three-stage model of initial binding followed by a reversible pore formation, cycling back and forth between partial insertion and subsequent full pore formation. [Fig. 4d].

### Nanobodies modify the dynamics of αSO pore formation

The toxicity of αSOs suggests that interfering with αSO mediate membrane permeabilization or perforation provides a therapeutic opportunity. Our single-particle approach enabled us to test whether ligands binding αSO affect the dynamics of pore formation. We discover that NB1 enhances pore formation and the translocation turnovers, suggesting a more flexible αSO structure and dynamic membrane interactions.

To achieve this, we tested two novel nanobodies which are, to our knowledge, the only NBs showing exclusive preference for αSO and not the monomer or fibril species [21]. First, we investigate the interaction between αSO and NBs by TEM. Binding of NB1 resulted in a blurred and less defined αSO structure while NB2 resulted in a population resembling free αSO [Fig. 5a and Supplementary Fig. 20]. Given the substantial changes observed from NB1 binding, we next evaluated the binding using our real-time single-vesicle recording assay as described in Fig 3. Fluorescently labeled αSO was added after 3 min by a continuous pumping system for 4 minutes at a low rate to allow membrane saturation followed by NB addition after 10 min (Fig 5c and Supplementary Fig. 21). NB1 addition resulted in a fluorescent quenching of the αSO, allowing recording of the binding rate of NB1 [Supplementary Fig. 22]. Surprisingly, no difference in binding rate is observed across liposome sizes or αSO densities [Supplementary Fig. 23]. In contrast, NB2 addition did not result in fluorescent quenching [Supplementary Fig. 22].

**Figure 5.**
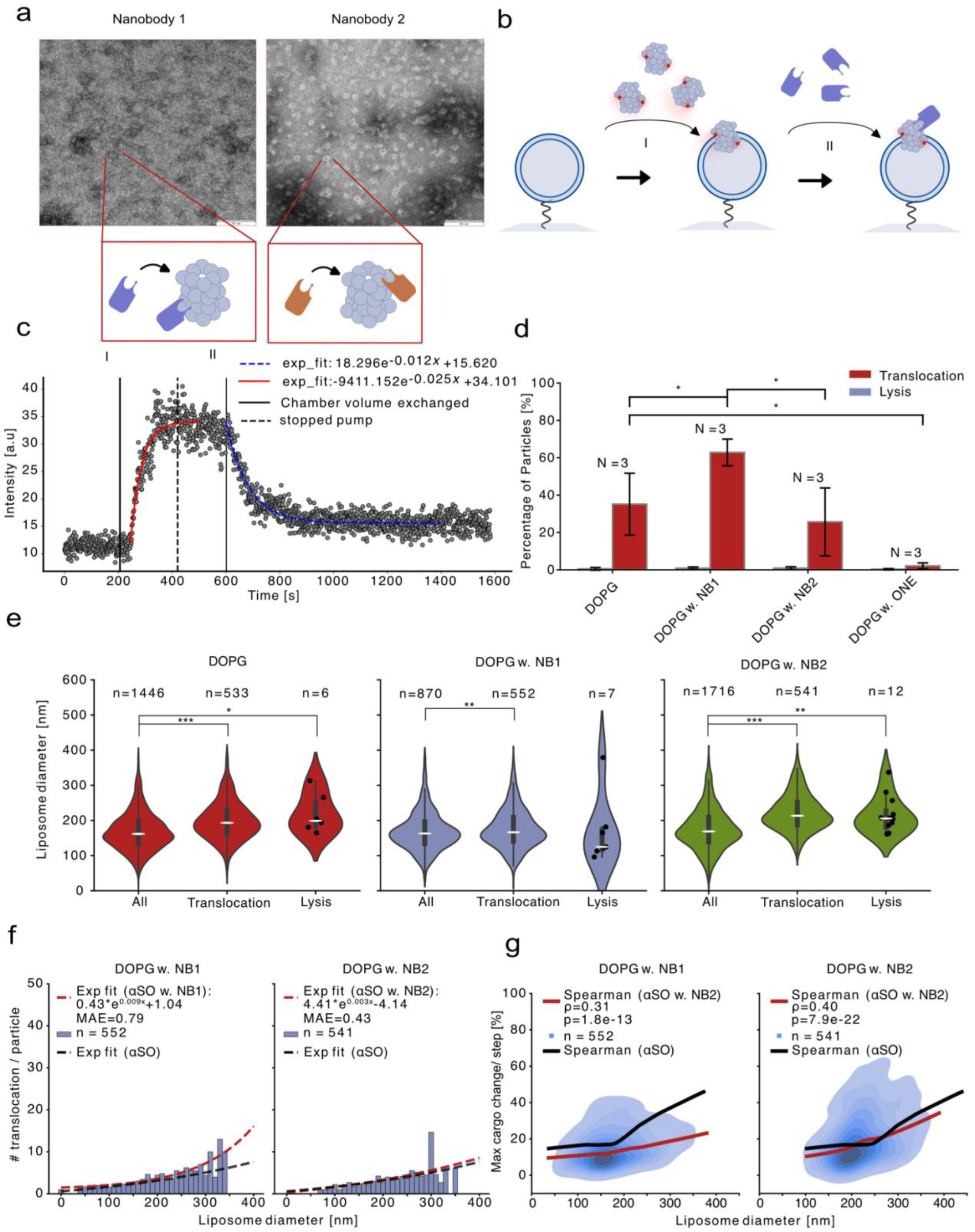
Nanobodies modulate the translocation of small molecules from the lumen. (a) nsTEM images confirming binding of NB1 and NB2 to αSO. (b) Schematic illustration of real-time imaging of nanobody binding to membrane-associated αSO. Fluorescently labeled αSOs are first injected into the microscope channel, followed by the injection of nanobodies. (c) representative time trajectory of real-time recording of αSO recruitment followed by NB1 binding. (d) statistics of αSO pore activity for DOPG vesicles with subsequent addition of either no nanobodies, NB1, or NB2, respectively. Additionally, we measured αSO pore activity for a chemically linked and stabilized αSO using ONE. Here the presence of NB1 results in a significant increase in activity and a stabilized ONE αSO shows significantly less activity. (e) Distributions of αSO pore activity for DOPG liposomes with no nanobodies (red), NB1 (blue) or NB2 (green) present, plotted against the liposome diameter. The distribution of pore-forming events in absence of nanobodies (red) is highly significantly shifted toward larger liposomes. The addition of both NB1 and NB2 additionally shows that pore-forming events remain highly significantly shifted toward larger liposomes. Asterisks indicate the following p-values with significance * = 0.05, ** = 0.01, *** = 0.005 based on a one-sided Mann-Whitney U test. (f) Quantification of the number of step-like translocations of dyes through the αSO pores. The presence of NB1 results in a more exponentially increasing distribution with liposome diameter than in the presence of NB2. (g) The quantity of lumen dyes translocated in a single translocation event. The presence of both NB1 and NB2 results in a positive correlation of % translocated with the liposome diameter.

To evaluate the influence of NB1 and NB2 binding to αSO we employed our real-time single-vesicle pore translocation recordings, as described in Fig. 3. Here we prepared 3 different oligomer preparations: αSO incubated with NB1 (αSO:NB1), αSO incubated with NB2 (αSO: NB2), and a 4-oxo-2-nonenal (ONE) chemical modified αSO (ONE-αSO), which stabilizes the structure, leading to a higher purification yield but also modifying the structure and membrane interactions of αSO [40]. The data were compared to the translocation in DOPG vesicles to allow statistical comparison. Interestingly, binding of NB1 resulted in a significant increase of pore formation, with 62.85 ± 7.15 % of vesicles translocating compared to 35.16 ± 16.55% in absence of NBs [Fig. 5d and supplementary Fig. 24]. The pore formation is decreased for vesicles with αSO:NB2 to 25.67 ± 18.17 % of the vesicles. The ONE-αSO led to highly significantly reduced pore formation with 2.10 ± 1.60 % translocating liposomes. We observed both NB1 and NB2 resulted in a significant upshift in curvature dependency for the pore-forming events (Fig. 5e) (P-value = 0.0054 and 8.32·10^−62^ MWU-test and P-value = 0.058 and 3.99·10^−58^ KS-test), although the shift for αSO:NB1 is markedly reduced, indicating that the effect of NB1 on pore formation is more significant in curved membranes.

Again, the real-time resolution allowed us to analyze the NB effect on translocation turnovers as a function of liposome size [Fig. 5f]. Here the fit from the absence of NB [from Fig 4b] is overlaid (black line) to compare the effects of the nanobodies (red line). NB1 changes the properties of αSO as the growth rate from the exponential fit is increased from 0.003 to 0.009. Thus, larger liposomes with less curved membranes are more susceptible to frequent translocation. In contrast, NB2 does not result in any detectable changes. Additionally, we quantified the quantity of the lumen dyes translocated in a single translocation event [Fig. 5g] compared to the absence of NB (black line). We find that NB1 results in a slightly decreased correlation between % translocated and liposome diameter with the Spearman coefficient decreasing from 0.36 to 0.31, as expected from the elevated translocation frequency. Again, NB2 does not result in any detectable changes. Taken together, NB1 results in a significantly increased pore formation and an elevated translocation turnover, suggesting a less stable or more flexible αSO structure, allowing for a more dynamic membrane interaction. The chemically modified ONE-αSO resulted in a significant decrease in pore-formation, again suggesting that a more rigid structure prevents the crucial dynamic αSO membrane interaction for pore formation.

## Discussion

Here, we developed a single-vesicle screening platform that enables a comprehensive biophysical model of αSO-membrane interactions for toxic αSO in unprecedented detail. We demonstrate real-time molecular translocation through αSO pores, visualized at single-vesicle resolution and employ single-channel electrical recordings to analyze pore formation in planar lipid bilayers. By integrating data from various methods, we propose a novel three-stage model for αSO-membrane interactions consisting of initial membrane recruitment followed by reversible cycling between partial insertion to full pore formation.

Our single-vesicle data reveal that initial recruitment is both lipid- and charge-dependent, with a notable curvature-promoted effect. Using real-time single-vesicle imaging, we elucidate how individual αSOs become fully integrated into membranes and function as pores, translocating small molecules across membranes. Negative membrane charge plays a key role in pore incorporation. Strikingly, while αSO recruitment favors curved membranes, pore formation predominantly occurs in less curved membranes. This suggests that recruitment is independent of charge-promoted reorientation and pore integration.

Our real-time single-vesicle recordings provide dynamic insights, revealing that individual liposomes undergo multiple translocations. The number of translocations positively correlates with liposome size, highlighting the highly dynamic nature of pore formation cycling between partial insertion to full pore formation. This process is modulated by membrane composition with pore formation being more pronounced and appearing to be more stable in anionic DOPG vesicles compared to anionic DOPS vesicles. Interestingly, pore formation can be influenced by oligomer-binding nanobodies. One of the nanobodies significantly increased pore formation and enhanced dynamics, suggesting an altered αSO structure that enhances membrane interaction.

Our results indicate that pore formation can be modulated by lipid composition and ligands, offering a biophysical understanding of αSO toxicity and providing a screening platform to identify ligands that alter αSO structure and potentially mitigate PD-related cellular dysfunction.

Few studies have investigated αSO membrane recruitment, often relying on bulk measurements or qualitative methods that fail to capture the inherent heterogeneity of the recruitment process. [23, 24]. To our knowledge, the only single vesicle study on αSO recruitment did not detect binding of αSO to the POPC membranes of giant unilamellar vesicles but clearly showed binding to DOPG and DOPS membranes [24]. However, a microfluidic bulk study detected αSO recruitment to DOPC membranes of SUVs and LUVs, consistent with our observations of αSO recruitment by DOPC membranes [23]. However, the previous study was limited to only two groups of vesicle sizes and provided only bulk data. Additionally, our setup enables real-time direct measurements, directly showcasing that the αSO is pore-forming.

The pore-forming ability of αSO has been suggested multiple times based on structural analysis [41, 42] and vesicle permeabilization studies [8, 24, 27, 41, 43]. Here, we provide evidence for pore formation using highly sensitive single-channel planar bilayer measurements, which is highly sensitive. A previous study demonstrated real-time pore formation in GUV vesicles [27], speculated to be driven by transient non-equilibrium processes. However, limited data prevents firm conclusions. It has also been shown that oligomer-membrane interactions can be inhibited[44], supporting our suggested three-stage model, as recruitment does not necessarily lead to pore formation.

Importantly, our assay enables high throughput screening of lipid compositions and curvatures and provides a direct and sensitive method for screening additives such as nanobodies. This could facilitate the identification of crucial ligands that bind to αSO to mitigate PD-related cellular dysfunction. More broadly, this assay establishes a framework to carry out comprehensive membrane-protein interaction analysis for protein aggregates in neurodegenerative disease. Tau protein aggregation, a hallmark of Alzheimer’s disease, has been suggested to permeabilize lysosomes[45], yet a detailed membrane-protein interaction analysis is lacking. The real-time single-vesicle platform enables screening at millisecond resolution up to days-long observations. Studies on the internalization of preformed fibrils have shown that α-synuclein aggregates perforate endolysosomes in neurons[46]. However, real-time single-vesicle analysis of synthetic membranes mimicking endolysosomes or isolated endolysosomes is lacking, which is essential for a comprehensive biophysical understanding. Such insights could pave the way for therapeutic interventions to prevent endolysosomal damage.

## Materials & Methods

### Materials

Unless otherwise stated all lipids and liposome extruder parts were acquired from Avanti Polar Lipids (Alabaster, Alabama, USA). Fluorophores including fluorophore-conjugated lipids were bought from ATTO-TEC GmbH (Siegen, Germany). Phosphate-buffered saline (PBS) was purchased from VWR (VWR International, Radnor, PA). For measurements containing CaCl_2_, PBS and the desired amount of CaCl_2_ stock were mixed to a final concentration of 1.2 mM CaCl_2_. All water was type 1 grade produced by MilliQ water purification system unless stated otherwise and all solvents.

### Expression and purification of α-Syn and αSO

Expression and purification of α-synuclein (α-Syn) followed the protocol described by Paslawski et al. [30]. Wild-type (WT) α-Syn was expressed in Escherichia coli BL21(DE3) cells containing the plasmid vector pET11a. Initially, cells were cultured on ampicillin-containing agar plates at 37°C overnight. Subsequently, they were transferred to an autoinduction medium and incubated at 37°C for 6 hours, followed by harvest via centrifugation at 4500 rpm and 4°C for 20 minutes. The resulting cell pellets were resuspended in osmotic shock buffer (30 mM Tris-HCl, 40% sucrose, 2 mM EDTA, pH 7.2) and subjected to centrifugation at 7000g and 20°C for 30 minutes. After collection, the pellets were dissolved in ice-cold water supplemented with 40 µL of saturated MgCl_2_ per 100 mL of medium while maintained on ice. The supernatant, obtained by centrifuging the dissolved pellets at 9000g and 4°C for 30 minutes, was then adjusted to pH 3.5 with HCl. After centrifugation at 9000g and 4°C for 20 minutes, the soluble proteins were collected in the supernatant, followed by adjustment of pH to 7.5 and storage at -80°C.

The solution was subsequently filtered through a 0.22 μm filter before being loaded onto a Q-Sepharose column (3×5 ml HiTrap Q HP) with a 20 mM Tris-HCl buffer at pH 7.5 for equilibration, followed by elution using a buffer containing 20 mM Tris-HCl and 1 M NaCl. Elution was carried out with a gradient of elution buffer ranging from 0% to 50%. Fractions containing α-Syn were collected and subjected to SDS-PAGE analysis to confirm protein purity [Supplementary Fig. 1a]. Pure fractions were pooled, dialyzed against Milli-Q (MQ) water, lyophilized, and stored at -20°C.

αSO were prepared by diluting lyophilized α-Syn to a concentration of 9-10 mg/mL in phosphate-buffered saline (PBS), followed by filtration using a 0.2 μm pore size syringe filter. The solution was then incubated in a Heating Shaking Dry bath (ThermoFisher Scientific) at 37°C for 3-5 hours. Prior to injection into the ÄKTA, the samples were centrifuged at 12000g for 5 minutes to remove large aggregates. After equilibration of a Superose 6 gel filtration column with PBS, αSO was eluted at a flow rate of 0.75 mL/min. Fractions containing αSO [Supplementary Fig. 1c], were pooled and concentrated using a 100 kDa Amicon Ultra-4 cutoff conical ultrafiltration unit (Merck Millipore Ltd., Tullagreen, Co., Cork, Ireland) at 4°C before storage at -20°C. Protein purity was reassessed using SDS-PAGE [Supplementary Fig. 1a]. All αSO concentrations are expressed in terms of mole of monomers.

### Expression and purification of nanobodies

A BL21 E. coli strain was transformed with a pET11d vector encoding either nanobody-1 (NB1) or nanobody-2 (NB2). Transfected E. coli cells were plated on agar plates containing Lysogeny broth (LB) medium supplemented with 2% glycerol and 1 mM MgCl^2^, followed by overnight incubation at 37°C. Colonies were then transferred to flasks containing LB medium supplemented with 0.1 mg/mL ampicillin and incubated for two hours at 37°C with agitation at 150 rpm using an Innova44 incubator shaker (New Brunswick, USA).

Induction of protein expression was initiated by adding 1 mM IPTG when the optical density (OD) of the culture reached 0.8, followed by further incubation for four hours at 37°C with agitation at 150 rpm. Subsequently, the cells were harvested by centrifugation at 4000 rpm for 20 minutes at 4°C, and the resulting pellets were resuspended in 50 mL of NB buffer (50 mM Tris pH 8, 150 mM NaCl) supplemented with a proteinase inhibitor tablet (Roche, 05 892 791 001) and DNase (Sigma, 9003-98-9).

After sonication (6 cycles of 20 seconds each), the suspension was centrifuged for 20 minutes at 10,000 g at 4°C, and the supernatant containing the expressed nanobodies was collected. The supernatant was then loaded onto a Histrap HP 1 mL column pre-equilibrated with NB buffer using a GE ÄKTA Pure system (ÄKTA, USA). Nanobodies were eluted from the column using a gradient of NB buffer containing 0.5 M imidazole. Fractions containing nanobodies were collected and desalted using a PD10 column. The purity of the nanobodies was assessed by SDS-PAGE [Supplementary Fig. 1a] To facilitate storage, 10% glycerol was added to the samples, which were then stored at -80°C.

### α-synuclein oligomer labeling

Labeling of αSO with ATTO 655 was done by mixing 150 μL of αSO (0.5-1 mg/mL), 50 μL of 1M bicarbonate, and 30 μL of 1.13 mM ATTO 655 NHS ester followed by two hours incubation on ice. Subsequently, the solution was centrifuged in a 100 kDa Amicon Ultra-4 cutoff conical ultrafiltration unit (Merck Millipore Ltd., Tullagreen, Co., Cork, Ireland) to get rid of the unconjugated dye. The labeled αSO was stored at -20°C. αSO-ATTO655 fluorescence was measured in the presence of NB on Cary Eclipse fluorescence spectrometer (Agilent Technologies, Santa Clara, CA) with excitation at 640 nm and emission at 655 and 850 nm. Spectra were recorded with slit widths of 10 nm (excitation) and 10 nm (emission) at 20°C.

### Liposome preparation

The preparation of liposomes was done according to an earlier described protocol [28, 47]. A series of unilamellar vesicles were prepared using 1,2-dioleoyl-sn-glycero-3-phospho-L-serine (DOPS), 1,2-Dioleoyl-sn-glycero-3-phosphocholine (DOPC), 1,2-Dioleoyl-sn-glycero-3-phosphoglycerol (DOPG), 0.5% 1,2-distearoyl-sn-glycero-3-phosphoethanolamine-N-[biotinyl(polyethylene glycol)-2000] (DSPE-PEG(2000) Biotin), and 0.5% ATTO488-1,2-Dioleoyl-sn-glycero-3-phosphoethanolamine (DOPE). The lipid composition ranged from 0% to 99% DOPG and 0% to 99% DOPS. The remaining lipids up to 100% were the neutral DOPC lipids. All lipid stocks were stored in chloroform at -20°C and mixed in glass vials. Chloroform was evaporated under a stream of N_2_, followed by an hour of incubation in a vacuum to form a lipid film.

Liposomes were formed by rehydrating the film in phosphate-buffered saline (PBS) (VWR International, Radnor, PA) containing 1.2 mM CaCl_2_ and allowed to self-assemble in the dark for 30 minutes.

The liposomes underwent ten cycles of flash-freezing and thawing to ensure unilamellar vesicles, followed by extrusion through a 200 nm Nuclepore Track-Etch membrane (GE Healthcare, Uppsala, Sweden) using an Avanti mini-extruder (Avanti Polar lipids Inc., Alabama, USA). Liposomes were stored at 4°C for a maximum of two weeks from preparation to imaging.

Liposomes with encapsulated molecules were made as described with the following additions to the protocol and described earlier [28, 37]. The rehydration buffer contained either 200 μM ATTO 488-dextran 4kDa (TdBLabs, Uppsala, Sweden) or 600 μM ATTO655 carboxy. In the case of ATTO 488-dextran 4kDa, ATTO488-DOPE was replaced with ATTO655.

### Acquisition of TIRF Microscopy data

Glass slides were cleaned by 10 minutes of sonication in 3x 2% Helmanex, 3x milliQ water, and 1x methanol. The cleaned glass surfaces were prepared using plasma-cleaned and activated pre-cleaned glass slices with attached Sticky-Slide VI 0.4 (Ibidi GmbH, Gräfelfing, Germany) and functionalized with PLL-g-PEG and PLL-g-PEG-biotin in a 100:1 ratio and incubated for 30 minutes. Excess PLL-g-PEG and PLL-g-PEG-biotin were removed by washing each well, followed by addition of a 0.1g/L neutravidin layer. Excess neutravidin was removed by washing each well with PBS.

Biotinylated liposomes were introduced into the system and allowed to immobilize, resulting in approximately 400 vesicles per field of view (FOV). Unbound liposomes were removed by washing with 3x chamber volumes of buffer.

A solution of 100 μL of 0.31 μM αSO-ATTO655 was added to the chamber and allowed to incubate for 10 minutes, after which unbound αSO were removed by washing 5 times with PBS containing 1.2 mM CaCl2. Imaging was conducted both before and after the addition of αSO using 7×7 images with 200 μm spacing between the centers of each FOV in an automated fashion. For real-time detection of encapsulated molecule leakage, time-lapse was made with a temporal resolution of 1 image/min in a single FOV for 5 hours. A Shenchen labv1 peristaltic flow pump (Baoding Shenchen Precision Pump Co., Ltd, Baodin, China) was integrated into the system for automatic αSO flow. Seven minutes into the acquisition, 0.3 mL of 0.31 μM αSO was infused into the system with a flow rate of 0.07 mL/min.

All single-particle experiments were performed using an Oxford Nanoimager S (Oxford NanoImaging, Oxford, UK), an inverted total internal reflection fluorescence microscope. Data were acquired using a 100x 1.41 NA oil-immersion objective at room temperature (19°C). Imaging utilized two solid-state lasers at 488 nm and 640 nm, and images were acquired with alternating lasers with laser powers set at 3.35% (<0.1 mW) and 5% (0.1 mW) and an exposure time of 200 ms. Image dimensions for each channel were 428 × 684 pixels with a dynamic range of 16-bit grayscale, recording 2 channels simultaneously. With a pixel size of 117 nm, the physical field of view (FOV) had dimensions of 50 μm x 80 per channel.

### TIRF real-time detection of NB binding

Glass slide preparation and liposome binding as described in the previous section. Images were acquired with a temporal resolution of 1 second for approx. 23 minutes. Images were acquired with alternating laser with an exposure time of 200 ms. After 3 minutes from start acquisition 0.3 mL 0.31 mM αSO was infused into the system. Subsequently, 10 minutes after start acquisition 0.3 mL 3.1 mM nanobody was infused into the system both were done with a flow rate of 0.07 mL/min by a peristaltic pump.

### Transmission electron microscopy (TEM)

A nanobody-binding antigen-binding fragment (NabFab) was conjugated to the nanobody in a ratio of 1:3. NB:NabFab was mixed with αSO in a 10:1 ratio.

Carbon-coated 400 mesh copper grids were glow discharged and 5 uL sample was added. The grids were washed with one drop of distilled water and stained with one drop of 1 % uranyl formate for negative staining (ns). The solution was blotted and dried. Electron microscopy was done on a Tecnai G2 Spirit (FEI company) and images were taken using a TemCam F416 camera (TVIPS). The sample protocol was used for both NB1 and NB2.

### Experimental Setup for Electrical Recordings of pore formation

To investigate pore insertion behaviors of αSO, we conducted electrical recordings using a setup involving vertical planar lipid membranes, following the established protocols [36, 48]. In summary, we employed either 1,2-diphytanoyl-sn-glycero-3-phosphocholine (DPhPC) or 1,2-dihexadecanoyl-sn-glycero-3-phosphoglycerol (DPhPG) to create a lipid bilayer on the Teflon film aperture within the chamber. The chamber was connected to a patch-clamp amplifier (Axopatch 200B, Axon Instruments) via two electrodes with an anode on the trans side and a cathode on the cis side. Each side of the chamber was filled with 1 mL of electrolyte buffer (1 M KCl, 50 mM Tris, pH 7.4), with or without the supplementation of 1.5 mM CaCl2. Varying concentrations of α-Syn monomer/oligomer were added to the trans side of the chamber and a voltage of +20 mV was applied for all the measurements. All recordings were conducted at a sampling frequency of 10 kHz and a room temperature of around 25 °C.

### Nanoparticle Tracking Analysis (NTA)

The hydrodynamic radius of the liposomes was determined using a NanoSight LM10 system (Malvern Instruments Ltd., Malvern, UK), fitted with a high sensitivity cCMOS camera (OrcaFlash2.8, Hamamatsu C11440, NanoSight Ltd) and a 405 nm laser. Each sample was diluted 1:1000 in PBS and measured in triplicates of 30-second recording with a camera level of 11 and a detection threshold of 3. Videos were recorded and analyzed using the NTA software (version 3.1, build 3.1.45). The ambient temperature was recorded manually and was approximately 20°C.

The relationship between the square root of the integrated signal from the membrane and liposome size is well-established, demonstrating a lognormal distribution of sizes as anticipated. By utilizing the average liposome size determined through Nanoparticle Tracking Analysis (NTA), it is possible to translate membrane intensity measurements into liposome sizes expressed in nanometers [35].

### Data analysis

#### Identification and co-localization software for TIRF multiplexing still imaging

The identification and co-localization of all liposomes and αSO were conducted using the membrane signal from the ATTO-488-DOPE membrane dye and co-localized to the αSO-ATTO-655 signal using in-house developed Python software, which was modified from recently published articles [28, 37, 47]. Each target was identified by using a Laplacian of Gaussian approximation, followed by selecting a region of interest (ROI) and a reference outer annulus to make an accurate local background correction. The software was designed to achieve nanometer-precise localization and colocalization across multiple imaging channels, effectively compensating for potential drifts caused by continuous fluid flow.

For each liposome formulation, the distribution of the colocalized αSO signal was square root transformed and fitted with a Gaussian distribution to obtain the mean and std. The data was normalized to the intensity before adding αSO which made us able to normalize to the overall shifts in liposome formulation.

### Identification and co-localization software for TIRF multiplexing real-time assay

Real-time measurements were saved as a stack of still images, where Identification and co-localization for each image in the stack were performed as described for the still imaging. After this the localized targets were connected through the z-stack using a Linear Assignment Problem Tracker (LAP Tracker), creating a time-resolved trajectory. This was done by using in-house developed Python software, which was modified from recently published articles [28, 37, 47].

By streamlining the tracking and analysis process, our software supports detailed investigations into the dynamics of liposome and αSO interactions, crucial for advancing our understanding of nanoscale biological processes.

### Statistical significance

We used the Mann-Whitney U test to compare the shift in mean since it does not assume normality and allows for heterogeneity of variance. A Kolmogorov–Smirnov test was used to compare whether two independent samples come from the same distribution, examining the differences in the entire distribution. The P-values for all statistical tests in the main text and the Supplementary Information are *P ≤ 0.05, **P ≤ 0.01, and ***P ≤ 0.001.

## Acknowledgments

This work was funded by Lundbeck Foundation grant No. R380-2021-1393 for M.G.M. Danish National Research Foundation grant no. 135 (CellPAT) for B.V.B, X.Z, and J.K. Lundbeck foundation grant No. R276-2018-671 and R453-2024-359 for J.N and D.E.O.

## Competing interests

The authors declare no competing interests.

## Supporting information

Is available for this paper.

## Data availability

Data supporting the findings of this study are available here https://anon.erda.au.dk/sharelink/F2mNpbWMlb. All raw data are available upon request.

## Code availability

Software code for treatment of single vesicle data and statistical analysis is available here: https://anon.erda.au.dk/sharelink/gnKqxEmisl

## Notes

### Competing Interest Statement

The authors have declared no competing interest.

### Summary of Updates

The author list was uploaded in a wrong order

## References

1. Auluck, P.K., G. Caraveo, and S. Lindquist, α-Synuclein: membrane interactions and toxicity in Parkinson’s disease. Annual review of cell and developmental biology, 2010. 26(1): p. 211–233.

2. Cookson, M.R., α-Synuclein and neuronal cell death. Molecular neurodegeneration, 2009. 4: p. 1–14.

3. Musteikytė, G., et al., Interactions of α-synuclein oligomers with lipid membranes. Biochimica et Biophysica Acta (BBA)-Biomembranes, 2021. 1863(4): p. 183536.

4. Mor, D.E., et al., Dynamic structural flexibility of α-synuclein. Neurobiology of disease, 2016. 88: p. 66–74.

5. Leverenz, J.B., et al., Proteomic identification of novel proteins in cortical lewy bodies. Brain Pathology, 2007. 17(2): p. 139–145.

6. Alam, P., et al., α-synuclein oligomers and fibrils: a spectrum of species, a spectrum of toxicities. Journal of neurochemistry, 2019. 150(5): p. 522–534.

7. Giehm, L., et al., Low-resolution structure of a vesicle disrupting α-synuclein oligomer that accumulates during fibrillation. Proceedings of the National Academy of Sciences, 2011. 108(8): p. 3246–3251.

8. Volles, M.J. and P.T. Lansbury, Vesicle permeabilization by protofibrillar α-synuclein is sensitive to Parkinson’s disease-linked mutations and occurs by a pore-like mechanism. Biochemistry, 2002. 41(14): p. 4595–4602.

9. van Rooijen, B.D., M.M. Claessens, and V. Subramaniam, Lipid bilayer disruption by oligomeric α-synuclein depends on bilayer charge and accessibility of the hydrophobic core. Biochimica et Biophysica Acta (BBA)-Biomembranes, 2009. 1788(6): p. 1271–1278.

10. Emin, D., et al., Small soluble α-synuclein aggregates are the toxic species in Parkinson’s disease. Nature communications, 2022. 13(1): p. 5512.

11. Awasthi, S., et al., Simultaneous determination of the size and shape of single α-synuclein oligomers in solution. ACS nano, 2023. 17(13): p. 12325–12335.

12. Roberts, H.L. and D.R. Brown, Seeking a mechanism for the toxicity of oligomeric α-synuclein. Biomolecules, 2015. 5(2): p. 282–305.

13. Lorenzen, N. and D.E. Otzen, Oligomers of α-synuclein: picking the culprit in the line-up. Essays in Biochemistry, 2014. 56: p. 137–148.

14. Chen, S.W., et al., Structural characterization of toxic oligomers that are kinetically trapped during α-synuclein fibril formation. Proceedings of the National Academy of Sciences, 2015. 112(16): p. E1994–E2003.

15. Lorenzen, N., et al., The role of stable α-synuclein oligomers in the molecular events underlying amyloid formation. Journal of the American Chemical Society, 2014. 136(10): p. 3859–3868.

16. Paslawski, W., et al., High stability and cooperative unfolding of α-synuclein oligomers. Biochemistry, 2014. 53(39): p. 6252–6263.

17. Asaadi, Y., et al., A comprehensive comparison between camelid nanobodies and single chain variable fragments. Biomarker Research, 2021. 9: p. 1–20.

18. Muyldermans, S., Nanobodies: natural single-domain antibodies. Annual review of biochemistry, 2013. 82: p. 775–797.

19. Caljon, G., et al., Using microdialysis to analyse the passage of monovalent nanobodies through the blood–brain barrier. British journal of pharmacology, 2012. 165(7): p. 2341–2353.

20. Muruganandam, A., et al., Selection of phage-displayed llama single-domain antibodies that transmigrate across human blood-brain barrier endothelium. The FASEB Journal, 2002. 16(2): p. 1–22.

21. Nielsen, J., et al., Nanobodies raised against the cytotoxic α-synuclein oligomer are specific for the oligomeric species and promote its cellular uptake. npj biosensing, 2025. In revision.

22. van Maarschalkerweerd, A., V. Vetri, and B. Vestergaard, Cholesterol facilitates interactions between α-synuclein oligomers and charge-neutral membranes. FEBS letters, 2015. 589(19): p. 2661–2667.

23. Greta Šneiderienė, M.A.C., Catherine K. Xu, Akhila Jayaram, View ORCID ProfileGeorg Krainer, View ORCID Profile William E. Arter, Quentin Peter, Marta Castellana-Cruz, Kadi L. Saar, Aviad Levin, Thomas Mueller, View ORCID Profile Sebastian Fiedler, Sean R. A. Devenish, Heike Fiegler, Janet R. Kumita, Tuomas P. J. Knowles, α-synuclein oligomers displace monomeric α-synuclein from lipid membranes. BioRxiv, 2023.

24. van Rooijen, B.D., M.M. Claessens, and V. Subramaniam, Membrane binding of oligomeric α-synuclein depends on bilayer charge and packing. FEBS letters, 2008. 582(27): p. 3788–3792.

25. Volles, M.J., et al., Vesicle permeabilization by protofibrillar α-synuclein: implications for the pathogenesis and treatment of Parkinson’s disease. Biochemistry, 2001. 40(26): p. 7812–7819.

26. Hannestad, J.K., et al., Single-vesicle imaging reveals lipid-selective and stepwise membrane disruption by monomeric α-synuclein. Proceedings of the National Academy of Sciences, 2020. 117(25): p. 14178–14186.

27. van Rooijen, B.D., M.M. Claessens, and V. Subramaniam, Membrane permeabilization by oligomeric α-synuclein: in search of the mechanism. PLoS One, 2010. 5(12): p. e14292.

28. Malle, M.G., et al., Single-particle combinatorial multiplexed liposome fusion mediated by DNA. Nature Chemistry, 2022. 14(5): p. 558–565.

29. Schmidt, S.G., et al., The dopamine transporter antiports potassium to increase the uptake of dopamine. Nature Communications, 2022. 13(1): p. 2446.

30. Paslawski, W., N. Lorenzen, and D.E. Otzen, Formation and characterization of α-synuclein oligomers. Protein Amyloid Aggregation: Methods and Protocols, 2016: p. 133–150.

31. Otzen, D.E., Antibodies and α-synuclein: What to target against Parkinson’s Disease? Biochimica et Biophysica Acta (BBA)-Proteins and Proteomics, 2024. 1872(2): p. 140943.

32. Nielsen, J., et al., Molecular properties and diagnostic potential of monoclonal antibodies targeting cytotoxic α-synuclein oligomers. npj Parkinson’s Disease, 2024. 10(1): p. 139.

33. Hatzakis, N.S., et al., How curved membranes recruit amphipathic helices and protein anchoring motifs. Nature chemical biology, 2009. 5(11): p. 835–841.

34. Kunding, A.H., et al., A fluorescence-based technique to construct size distributions from single-object measurements: application to the extrusion of lipid vesicles. Biophysical Journal, 2008. 95(3): p. 1176–1188.

35. Lohr, C., et al., Constructing size distributions of liposomes from single-object fluorescence measurements. Methods in enzymology, 2009. 465: p. 143–160.

36. Zhang, X., et al., Specific Detection of Proteins by a Nanobody-Functionalized Nanopore Sensor. ACS nano, 2023. 17(10): p. 9167–9177.

37. Zhang, X., et al., Deconvoluting the Effect of Cell-Penetrating Peptides for Enhanced and Controlled Insertion of Large-Scale DNA Nanopores. ACS Applied Materials & Interfaces, 2024. 16(15): p. 18422–18433.

38. Quist, A., et al., Amyloid ion channels: a common structural link for protein-misfolding disease. Proceedings of the National Academy of Sciences, 2005. 102(30): p. 10427–10432.

39. Lim, L. and M. Wenk, Neuronal membrane lipids–their role in the synaptic vesicle cycle, in Handbook of neurochemistry and molecular neurobiology. 2009.

40. Andersen, C., et al., Lipid peroxidation products HNE and ONE promote and stabilize alpha-synuclein oligomers by chemical modifications. Biochemistry, 2021. 60(47): p. 3644–3658.

41. Lashuel, H.A., et al., α-Synuclein, especially the Parkinson’s disease-associated mutants, forms pore-like annular and tubular protofibrils. Journal of molecular biology, 2002. 322(5): p. 1089–1102.

42. Fusco, G., et al., Structural basis of membrane disruption and cellular toxicity by α-synuclein oligomers. Science, 2017. 358(6369): p. 1440–1443.

43. Ghio, S., et al., Cardiolipin promotes pore-forming activity of alpha-synuclein oligomers in mitochondrial membranes. ACS chemical neuroscience, 2019. 10(8): p. 3815–3829.

44. Lorenzen, N., et al., How Epigallocatechin Gallate Can Inhibit α-Synuclein Oligomer Toxicity in Vitro♦. Journal of Biological Chemistry, 2014. 289(31): p. 21299–21310.

45. Rose, K., et al., Tau fibrils induce nanoscale membrane damage and nucleate cytosolic tau at lysosomes. Proceedings of the National Academy of Sciences, 2024. 121(22): p. e2315690121.

46. Sanyal, A., et al., Neuronal constitutive endolysosomal perforations enable α-synuclein aggregation by internalized PFFs. Journal of Cell Biology, 2024. 224(2): p. e202401136.

47. Malle, M.G., et al., Programmable RNA Loading of Extracellular Vesicles with Toehold-Release Purification. Journal of the American Chemical Society, 2024. 146(18): p. 12410–12422.

48. Maglia, G., et al., Analysis of single nucleic acid molecules with protein nanopores, in Methods in enzymology. 2010, Elsevier. p. 591–623.

