## Supplementary information for "Single-vesicle tracking of α-synuclein oligomers reveals pore formation by a three-stage model modulated by charge, curvature, lipids and ligands"

***Affiliation:***
[a] Interdisciplinary Nanoscience Center (iNANO), Aarhus University, Gustav Wieds Vej 14, 8000 Aarhus C, Denmark
[b]Department of Molecular Biology and Genetics, Aarhus University, Universitetsbyen 81, 8000 Aarhus C, Denmark

### Table of Contents

SI Fig. 1: Purification of nanobodies and alpha-synuclein. 3
SI Fig. 2: Representative example of the nanoparticle tracking analysis (NTA) 4
SI Fig. 3: Representative example of the size distribution of liposomes before and after the addition of αSO. 5
SI Fig. 4: Representative example of the size distribution of liposomes before and after the addition of αSO. 6
SI Fig. 5: Representative example of intensity shift in 640 channels upon addition of αSO. 7
SI Fig. 6: Representative example of intensity shift in 640 channels upon addition of αSO. 8
SI Fig. 7: Surface coating shows no unspecific binding of αSO. 9
SI Fig. 8: Quantification of the charge and curvature dependency of αSO recruitment of liposome membranes. 10
SI Fig. 9: Quantification of the charge and curvature dependency of αSO recruitment of liposome membranes. 11
SI Fig. 10: Extracted β-values from density quantification fit. 12
SI Fig. 11: Representative examples of real-time single-vesicle recordings demonstrating αSO pore formation and the translocation of small molecules. 12
SI Fig. 12: Representative examples of real-time single-vesicle recordings demonstrating no translocation. 13
SI Fig. 13: Representative examples of real-time single-vesicle recordings with αSM demonstrating limited translocation. 16
SI Fig. 14: Representative examples of real-time single-vesicle recordings with buffer only demonstrating no translocation. 16
SI Fig. 15: Representative current trajectories showing the pore formation of α-syn oligomer on neutral-charged planar lipid bilayer. 17
SI Fig. 16: Representative current trajectories showing the pore formation of α-syn oligomer on neutral-charged planar lipid bilayer in the presence of CaCl2. 19
SI Fig. 17: Representative current trajectories showing the pore formation of α-syn oligomer on the negatively charged planar lipid bilayer. 18
SI Fig. 18: Representative current trajectories showing the pore formation of α-syn oligomer on the negatively charged planar lipid bilayer in the presence of CaCl2 20
SI Fig. 19: Control experiment showing that α-syn monomer has no pore formation activity on the negatively charged planar lipid bilayer 20
SI Fig. 20: Encapsulation of ATTO-655 carboxy in DOPC, DOPG and DOPS containing liposomes. 21
SI Fig. 21: Representative of real-time trajectories of αSO binding followed by NB1 binding to single-vesicles. 22
SI Fig. 22: Nanobody 1 quenches ATTO655 signal. 23
SI Fig. 23: Quantification of αSO recruitment and nanobody binding. 24
SI Fig. 24: Representative examples of real-time single-vesicle recordings demonstrating αSO pore formation in presence of NBs. 25

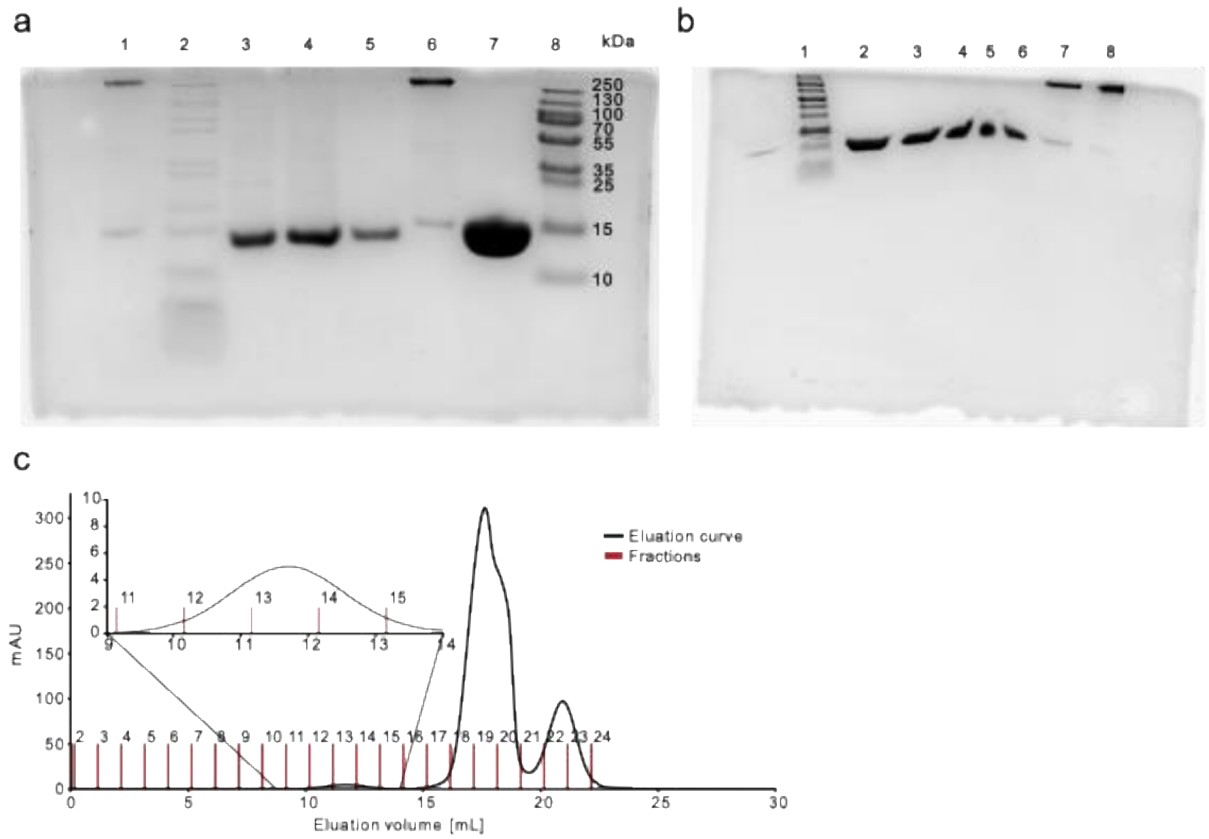


SI Fig. 1: Purification of nanobodies and alpha-synuclein. *a) SDS-PAGE of pooled protein fractions. Lane 1 + 6; αSO, lane 2; molecular weight marker proteins using Mark 12 unstained standard kit, lane 3+4; nanobody 1, lane 5; nanobody 2, lane 7; αSM, and lane 8; PageRuler™ Plus Prestained Protein Ladder. αSM and αSO were eluted from a Superose 6 gel filtration column and the nanobodies were eluted from a Histrap HP column. b) SDS-PAGE used for concentration determination of αSO. Lane 1; mark 12 ladder, Lane 2-6 increasing αSM concentration, lane 7+8; two αSO purifications. c) GE ÄKTA Pure system elution curve showing fractions (numbers 2-24) from αSO purification. The zoom-in (fractions 12-14) shows pooled fractions containing αSO. Fractions 18-20 contain αSM.*

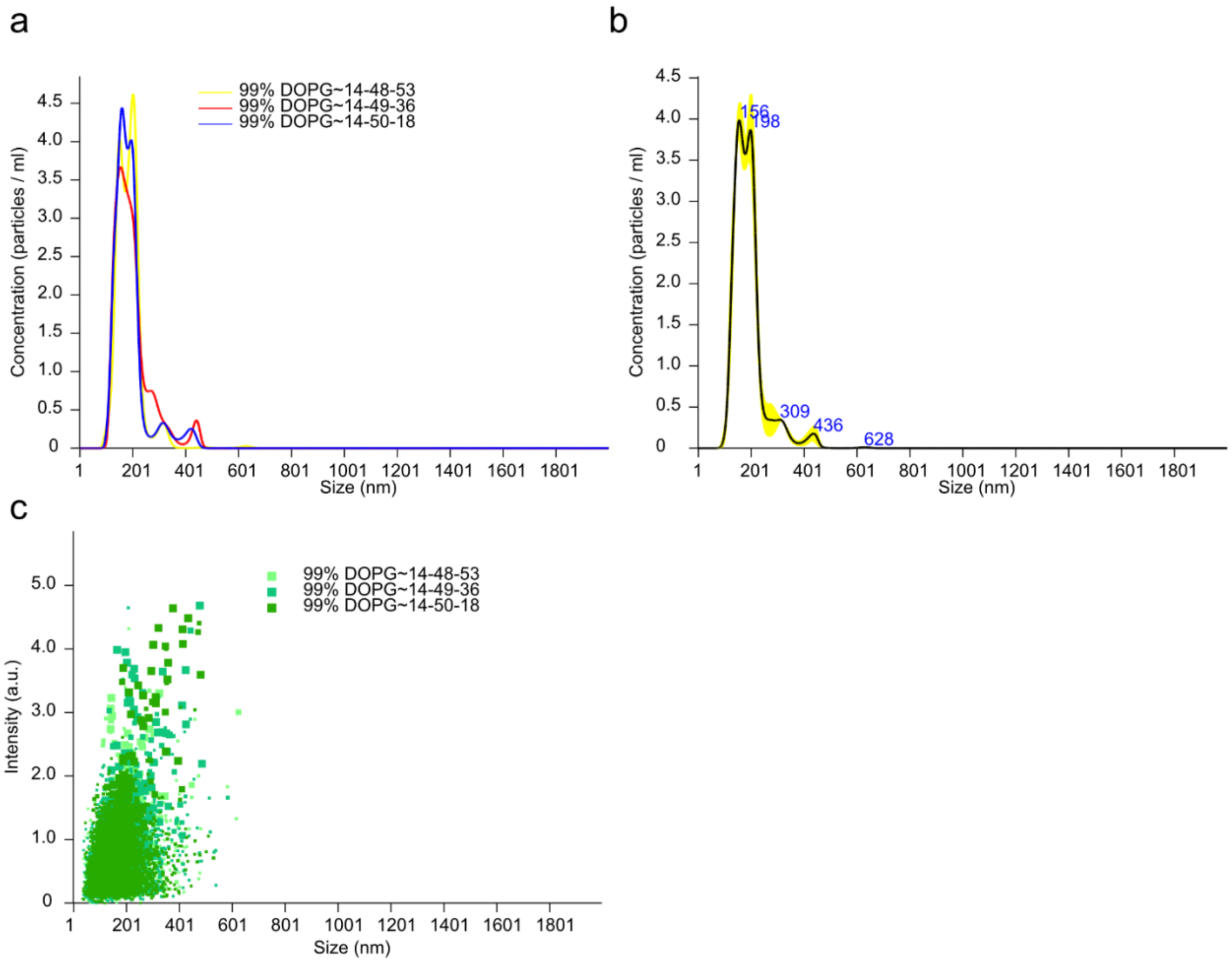


*SI Fig. 2: Representative example of the nanoparticle tracking analysis (NTA)*. Liposomes were injected into the NTA chamber and measured a) in triplicates of 30-second recording with a camera level of 11 and a detection threshold of 3. b) Triplicates were averaged to give a size distribution. c) Showing the intensity/size graph for all single liposomes measured.

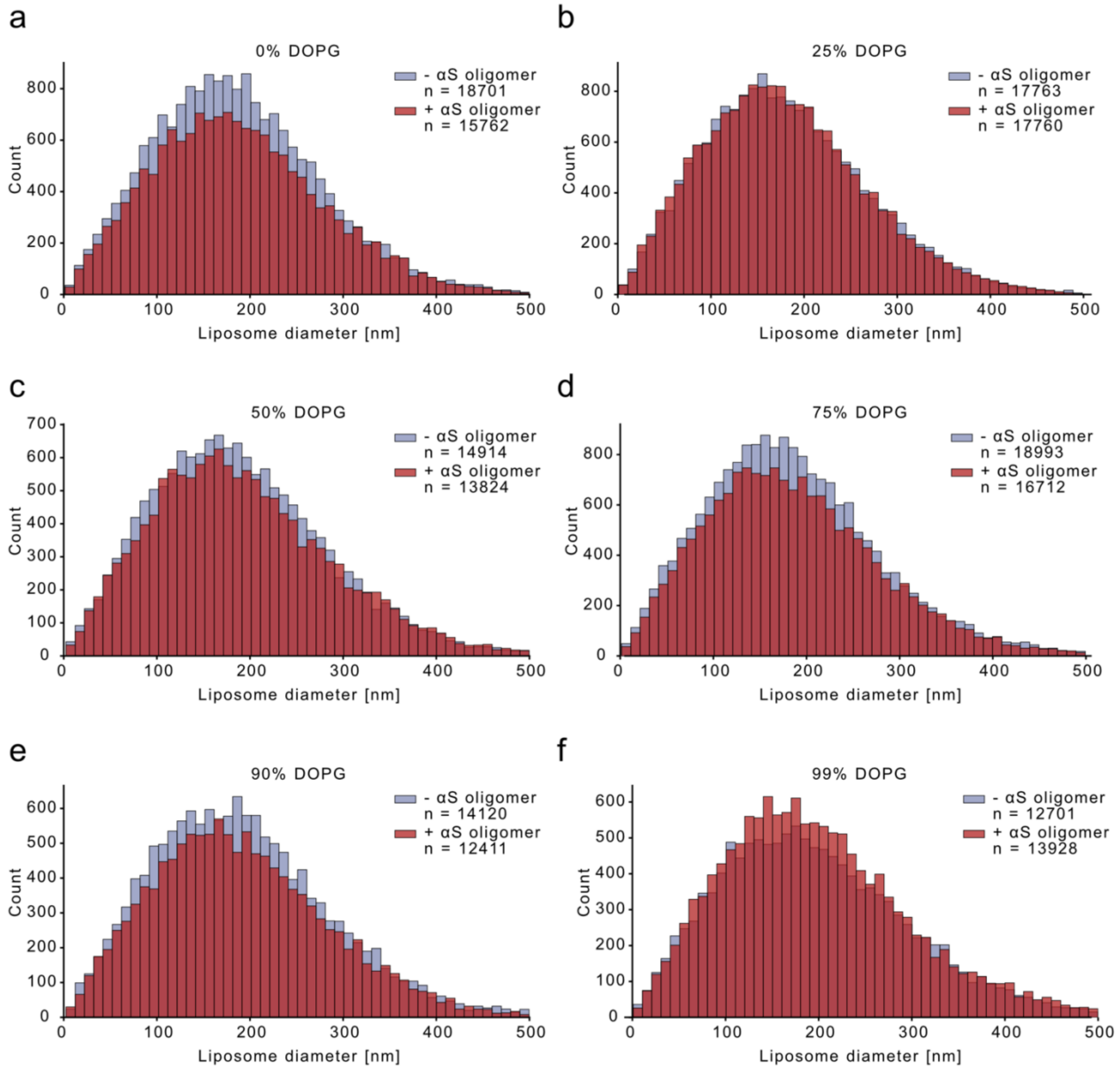


SI Fig. 3: Representative example of the size distribution of liposomes before and after the addition of αSO. *The intensities of liposomes from 48 randomly selected field-of-views (FOWs) were extracted before addition (blue). Another 48 random FOWs were selected after the addition of αSO (red) and a Kolmogorov–Smirnov test was used to determine if the two distributions were pulled from the same underlying distribution. A series of liposomes were tested where the membrane consisted of 0.5%DSPE-PEG(2000) Biotin, 0.5% ATTO488-DOPE and the rest a mix of DOPC and DOPG. Here the content was: a) 0% DOPG liposomes; p-value: 0.37. b) 25% DOPG liposomes; p-value: 0.75. c) 50% DOPG liposomes; p-value: 0.10. d) 75% DOPG liposomes; p-value: 0.33. e) 90% DOPG liposomes; p-value: 0.83. f) 99% DOPG liposomes; p-value: 0.08.*

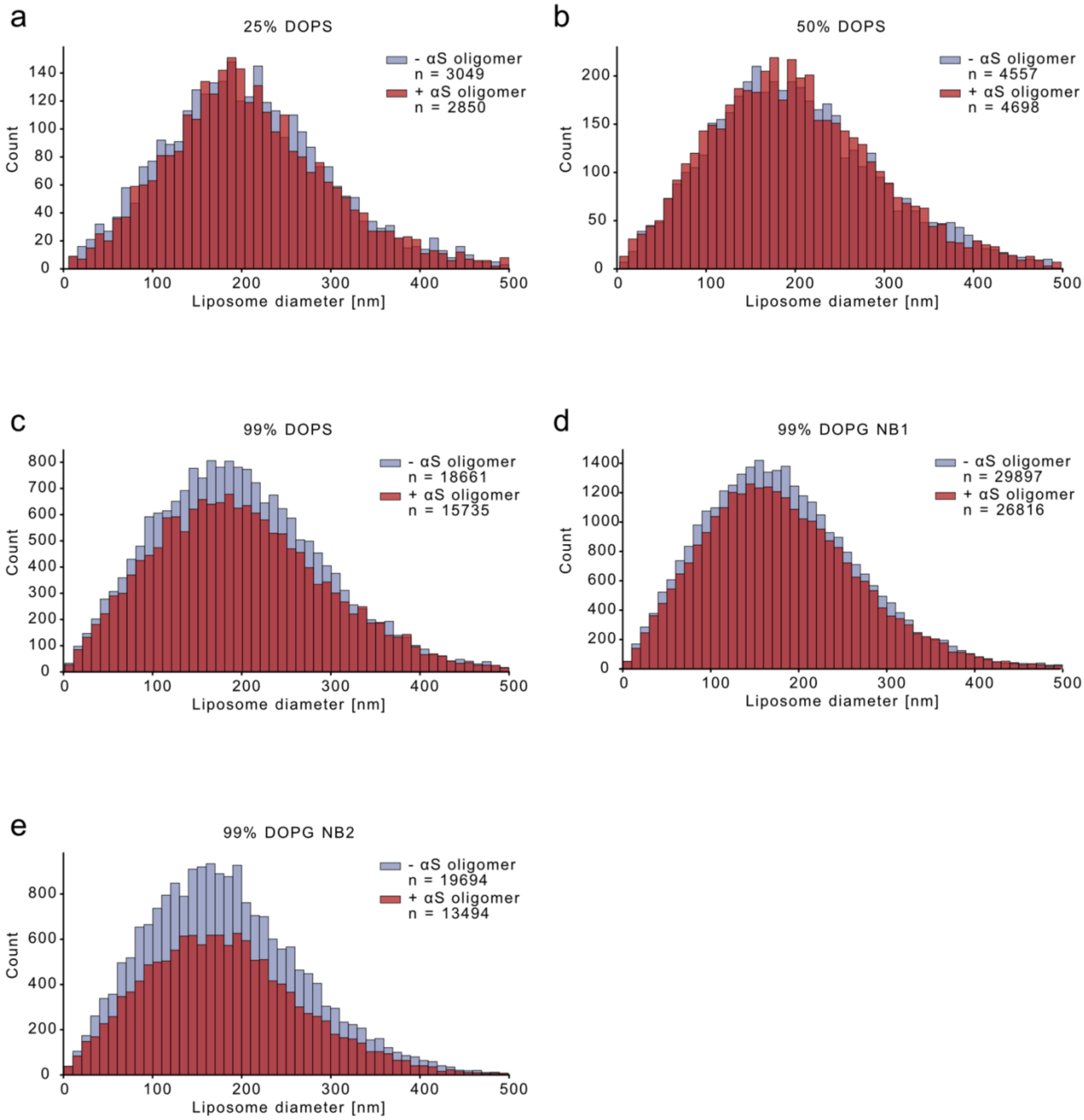


SI Fig. 4: Representative example of the size distribution of liposomes before and after the addition of αSO. *The intensities of liposomes from 48 randomly selected field-of-views (FOWs) were extracted before addition (blue). Another 48 random FOWs were selected after the addition of αSO (red) and a Kolmogorov–Smirnov test was used to determine if the two distributions were pulled from the same underlying distribution. A series of liposomes were tested where the membrane consisted of 0.5%DSPE-PEG(2000) Biotin, 0.5% ATTO488-DOPE and the rest a mix of DOPC and DOPS or DOPG. Here the content was: a) 25% DOPS liposomes; p-value: 0.53. b) 50% DOPS liposomes; p-value: 0.68. c) 99% DOPG liposomes; p-value: 0.32. d) 99% DOPG liposomes with NB1; p-value: 0.80. e) 99% DOPG liposomes with NB2; p-value: 0.47.*

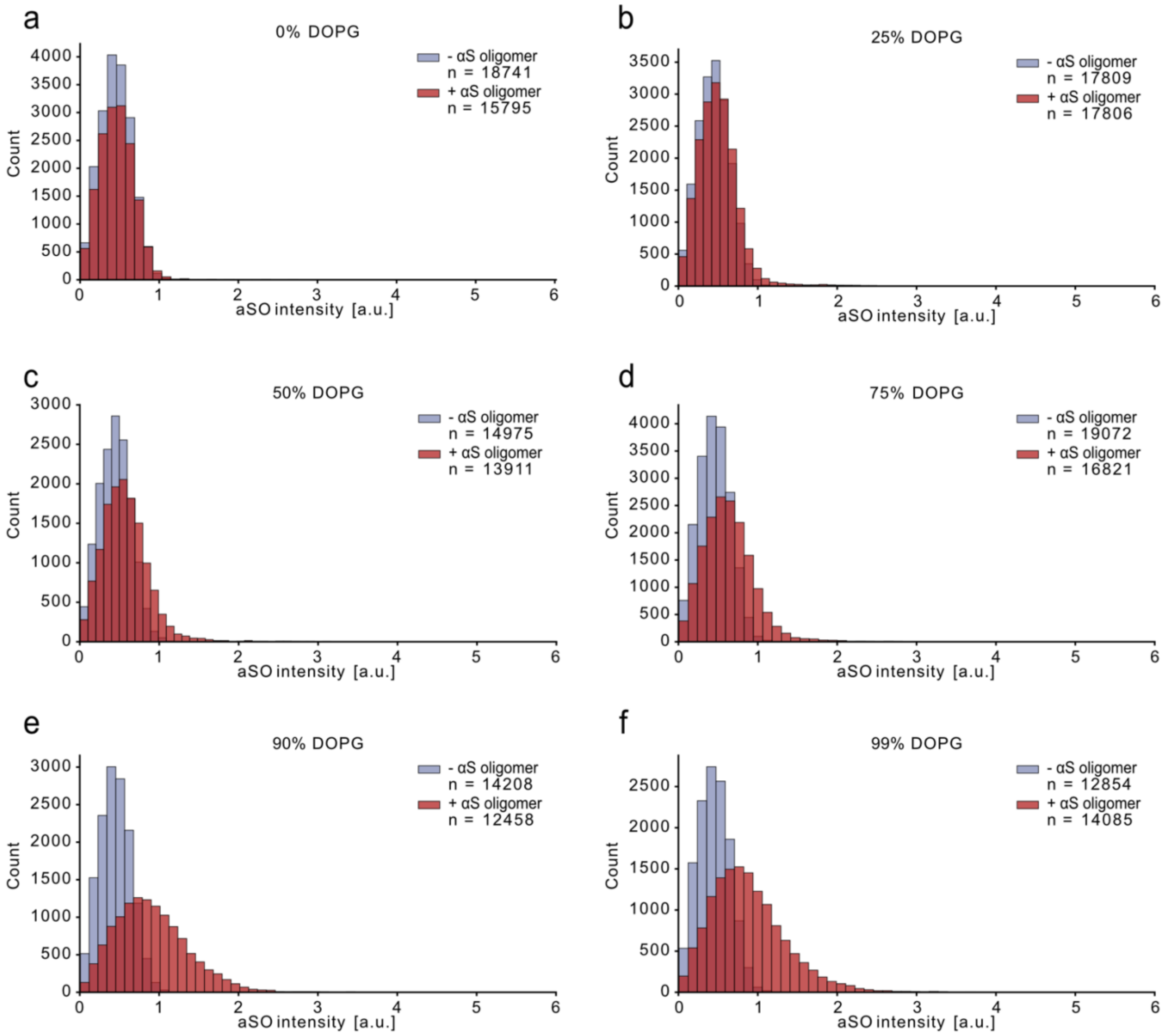


SI Fig. 5: Representative example of intensity shift in 640 channels upon addition of αSO. *The intensities from the co-localized spots in the liposome channel [Supplementary Fig. 4] were extracted before addition (blue). And again after the addition of αSO (red) a Kolmogorov–Smirnov test was used to determine if the two distributions were pulled from the same underlying distribution. a) 0% DOPG liposomes; p-value: 2.77e-07. b) 25% DOPG liposomes; p-value: 1.02e-52. c) 50% DOPG liposomes; p-value: 9.30e-257. d) 75% DOPG liposomes; p-value: 0.00. e) 90% DOPG liposomes; p-value: 0.83. f) 99% DOPG liposomes; p-value: 0.00.*

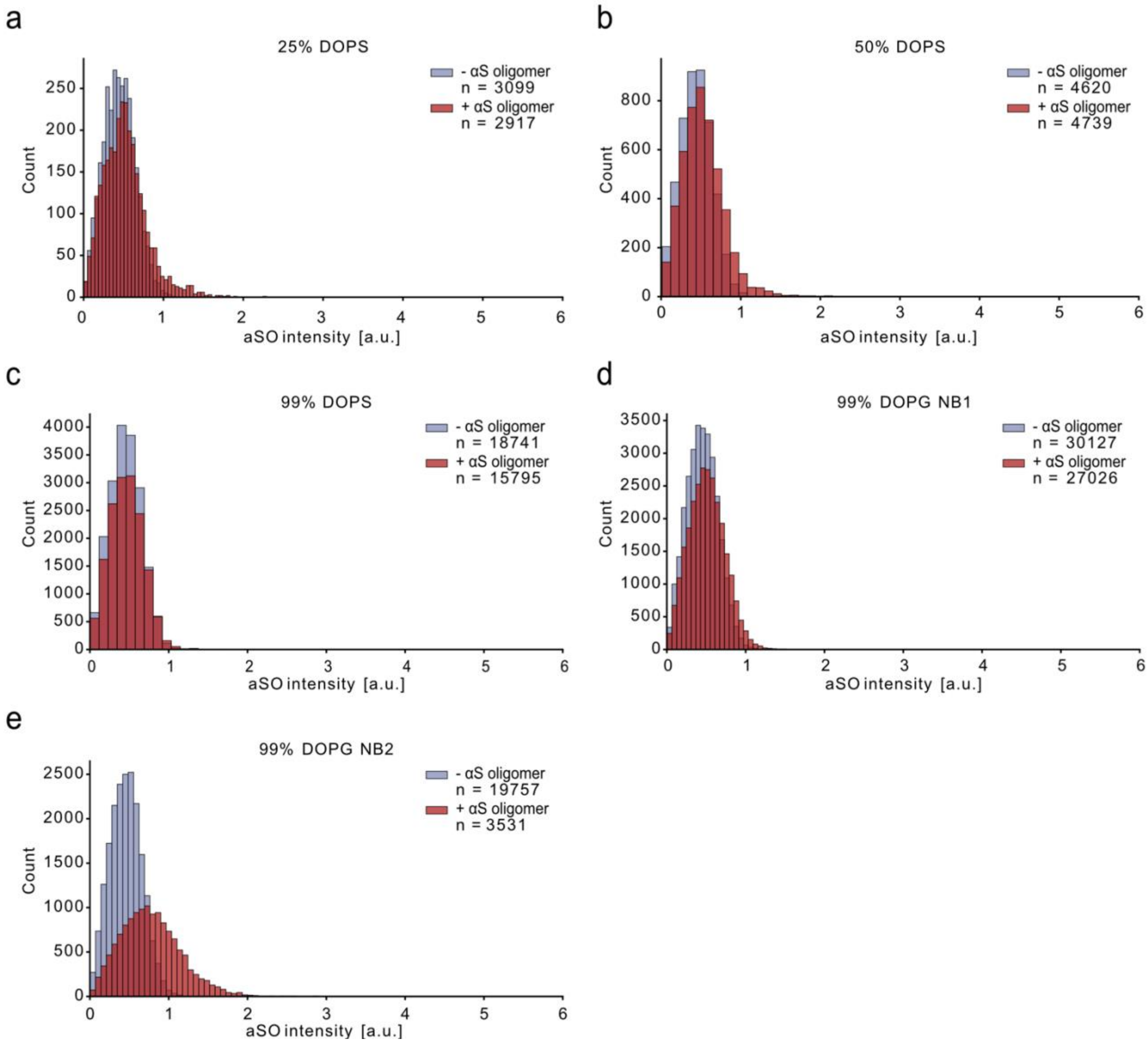


SI Fig. 6: Representative example of intensity shift in 640 channels upon addition of αSO. *The intensities from the co-localized spots in the liposome channel [Supplementary Fig. 4] were extracted before addition (blue). And again, after the addition of αSO (red) a Kolmogorov–Smirnov test was used to determine if the two distributions were pulled from the same underlying distribution. a) 25% DOPS liposomes; p-value:* 3.33e-16. *b) 50% DOPS liposomes; p-value:* 3.33e-15. *c) 99% DOPG liposomes; p-value: 2.77e-07. d) 99% DOPG liposomes with NB1; p-value:* 1.87e-146. *e) 99% DOPG liposomes with NB2; p-value: 0.00.*

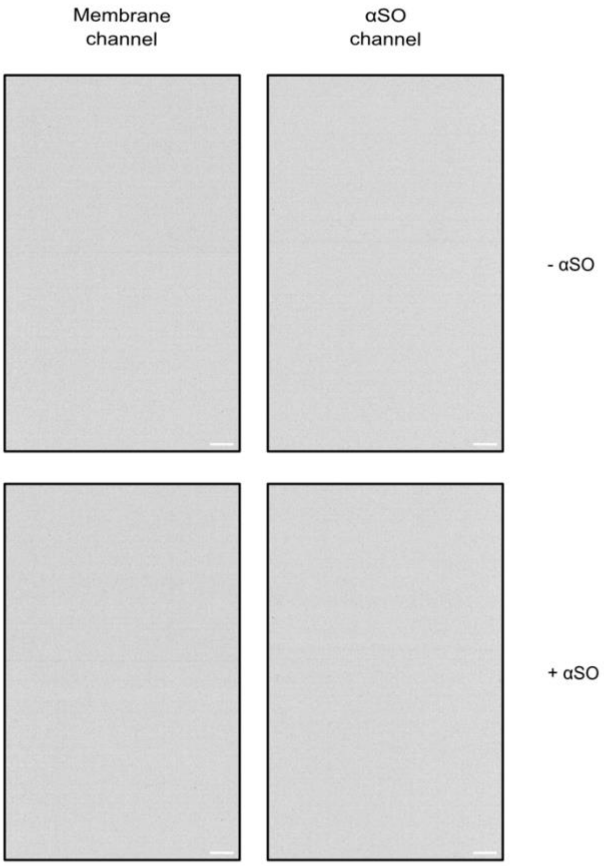


SI Fig. 7: Surface coating shows no unspecific binding of αSO. *Representative examples of images from the membrane channel (left) and the αSO channel (right) where no liposomes were tethered to the surface.*

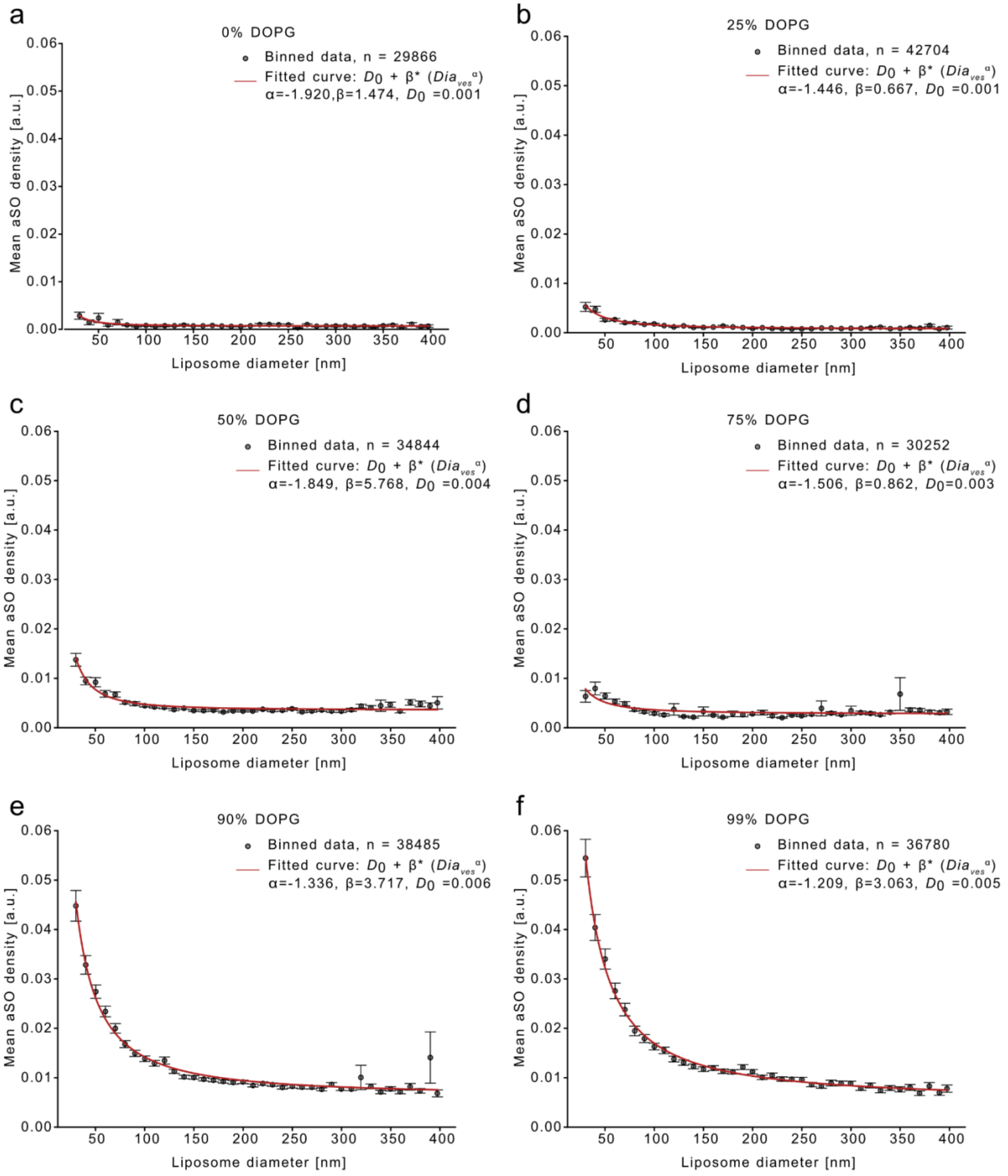


SI Fig. 3: Quantification of the charge and curvature dependency of αSO recruitment of liposome membranes. *The αSO density ($D_{\alpha SO}$) was calculated for all liposomes and binned into 10nm intervals. Each distribution was fitted with an offset power function:* $D_{\alpha SO} = D_0 + \beta * Dia_{liposome}{}^{\alpha}$. *The series of liposomes were tested where the membrane consisted of 0.5%DSPE-PEG(2000) Biotin, 0.5% ATTO488-DOPE, and the rest a mix of DOPC and DOPG. Here the content was: a) 0% DOPG b) 25% DOPG c) 50% DOPG d) 75% DOPG e) 90% DOPG f) 99% DOPG.*

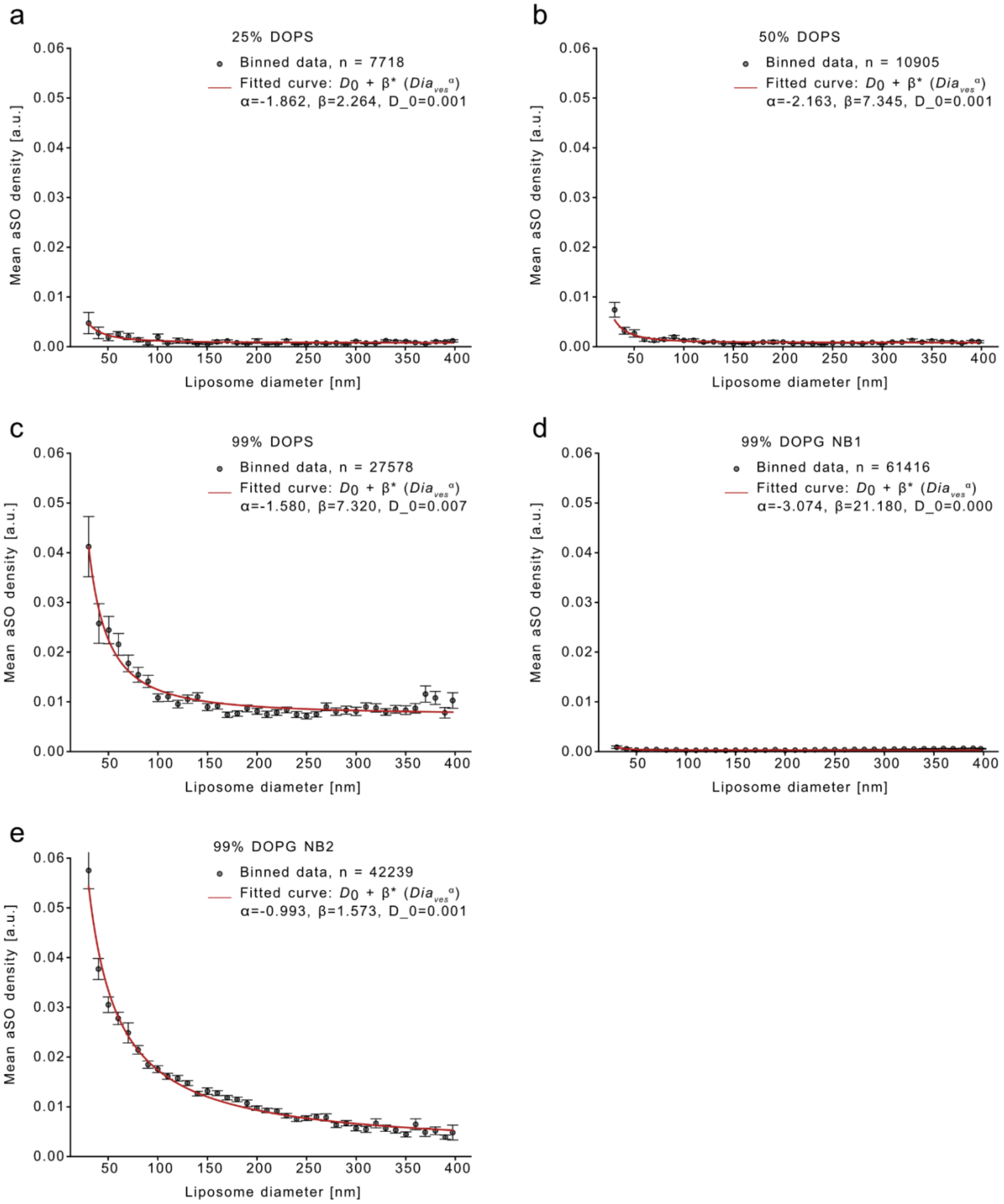


SI Fig. 4: Quantification of the charge and curvature dependency of αSO recruitment of liposome membranes. *The αSO density ($D_{\alpha SO}$) was calculated for all liposomes and binned into 10nm intervals. Each distribution was fitted with an offset power function:* $D_{\alpha SO} = D_0 + \beta * Dia_{liposome}{}^{\alpha}$. *The series of liposomes were tested where the membrane consisted of 0.5%DSPE-PEG(2000) Biotin, 0.5% ATTO488-DOPE and the rest a mix of DOPC and DOPS or DOPG. Here the content was: a) 25% DOPS b) 50% DOPS c) 99% DOPS d) 99% DOPG with NB1 e) 99% DOPG with NB2.*

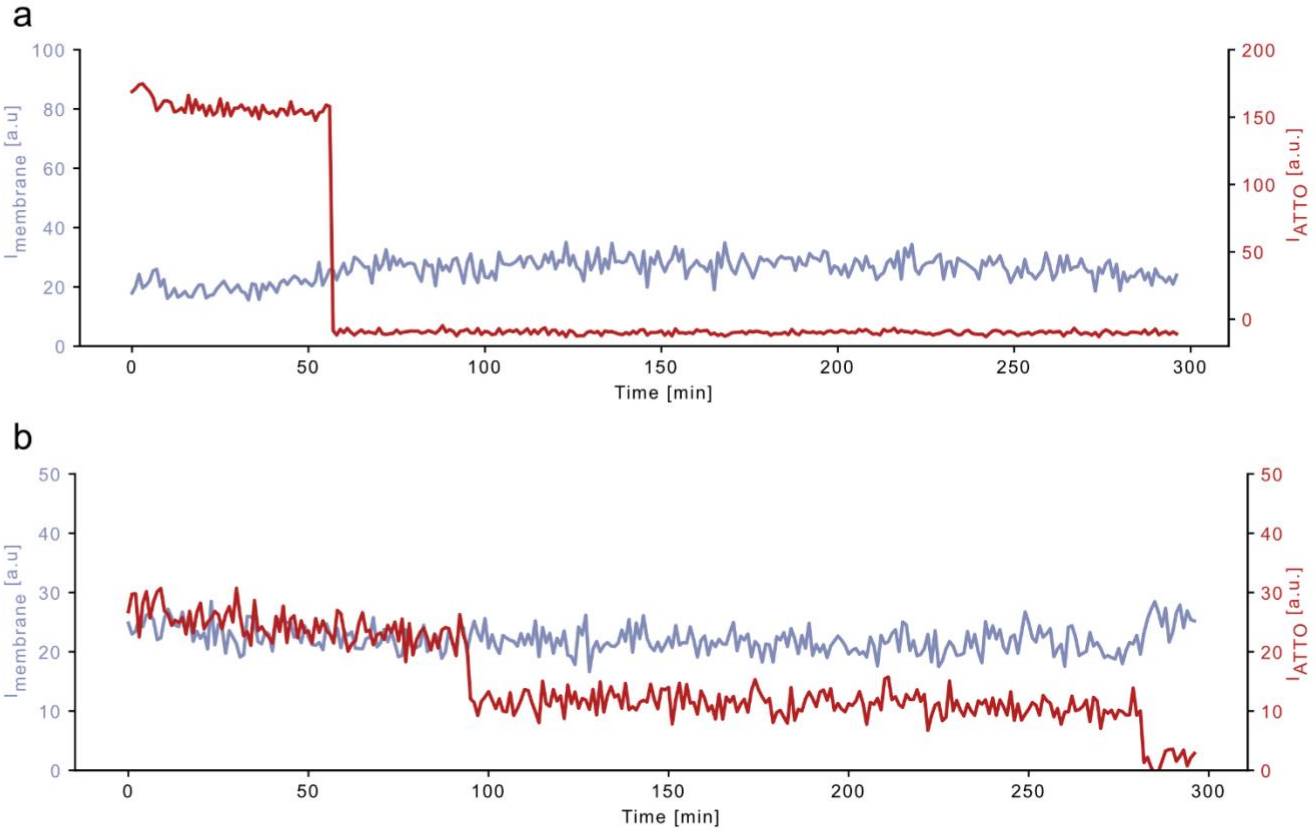


SI Fig. 10: Representative examples of real-time single-vesicle recordings demonstrating αSO pore formation and the translocation of small molecules. *99% DOPG liposomes with encapsulated ATTO655-carboxy, were monitored with a framerate of 1 image/min for 300 minutes. αSO were washed in the chamber after 7 minutes. Showing no membrane rupture (blue) for the two particles with a) one translocation event and b) two translocation events. Both trajectories show a stable liposome membrane and a dynamic pore formation with two distinct open pore formations allowing for dyes' translocation. The membrane signal and the ATTO655 signal are given in arbitrary units.*

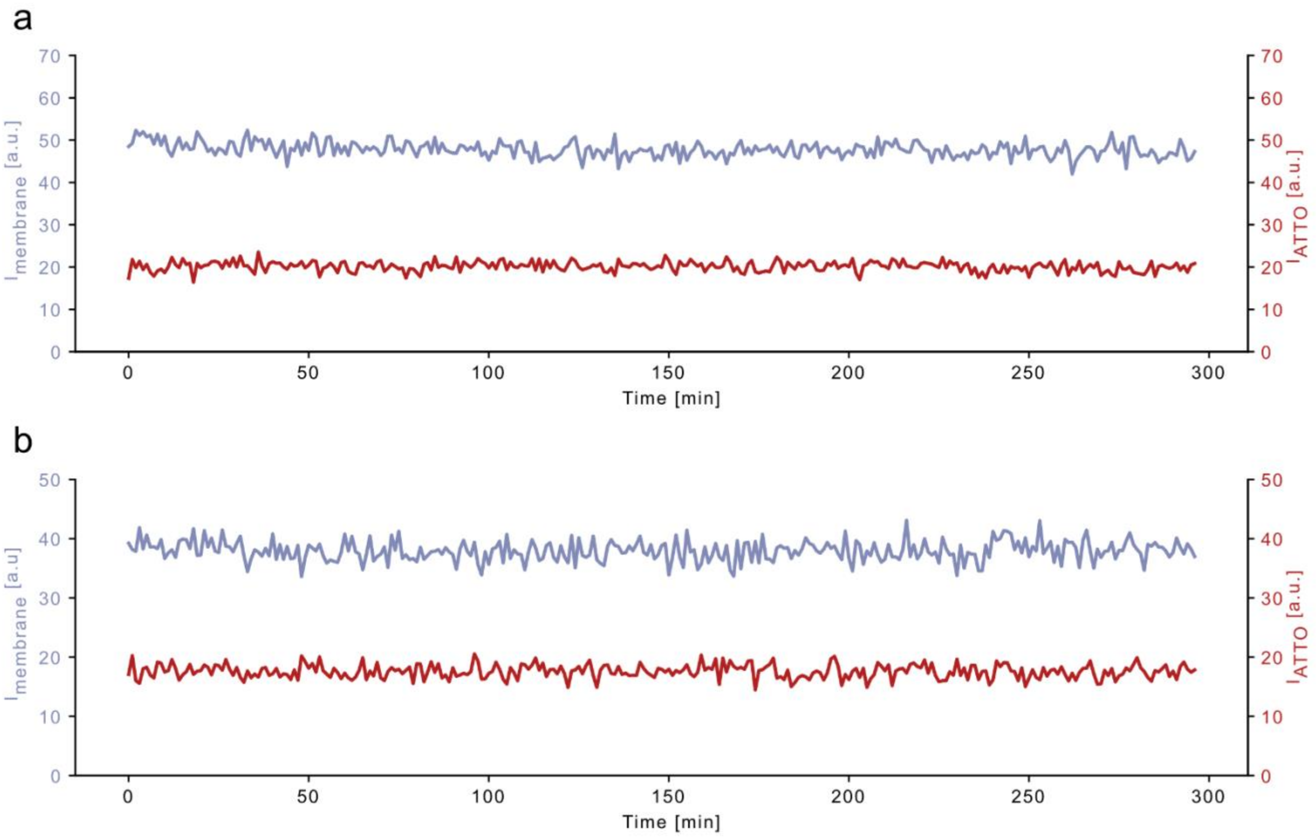


SI Fig. 11: Representative examples of real-time single-vesicle recordings demonstrating no translocation. *99% DOPG liposomes with encapsulated ATTO 488-dextran 4kDa, were monitored with a framerate of 1 image/min for 300 minutes. αSO were washed in the chamber after 7 minutes. Showing no membrane rupture (blue) for the two particles with a+b) no translocation events. Both trajectories show a stable liposome membrane but do not allow for dyes' translocation. The membrane signal and the ATTO 488-dextran 4kDa signal are given in arbitrary units.*

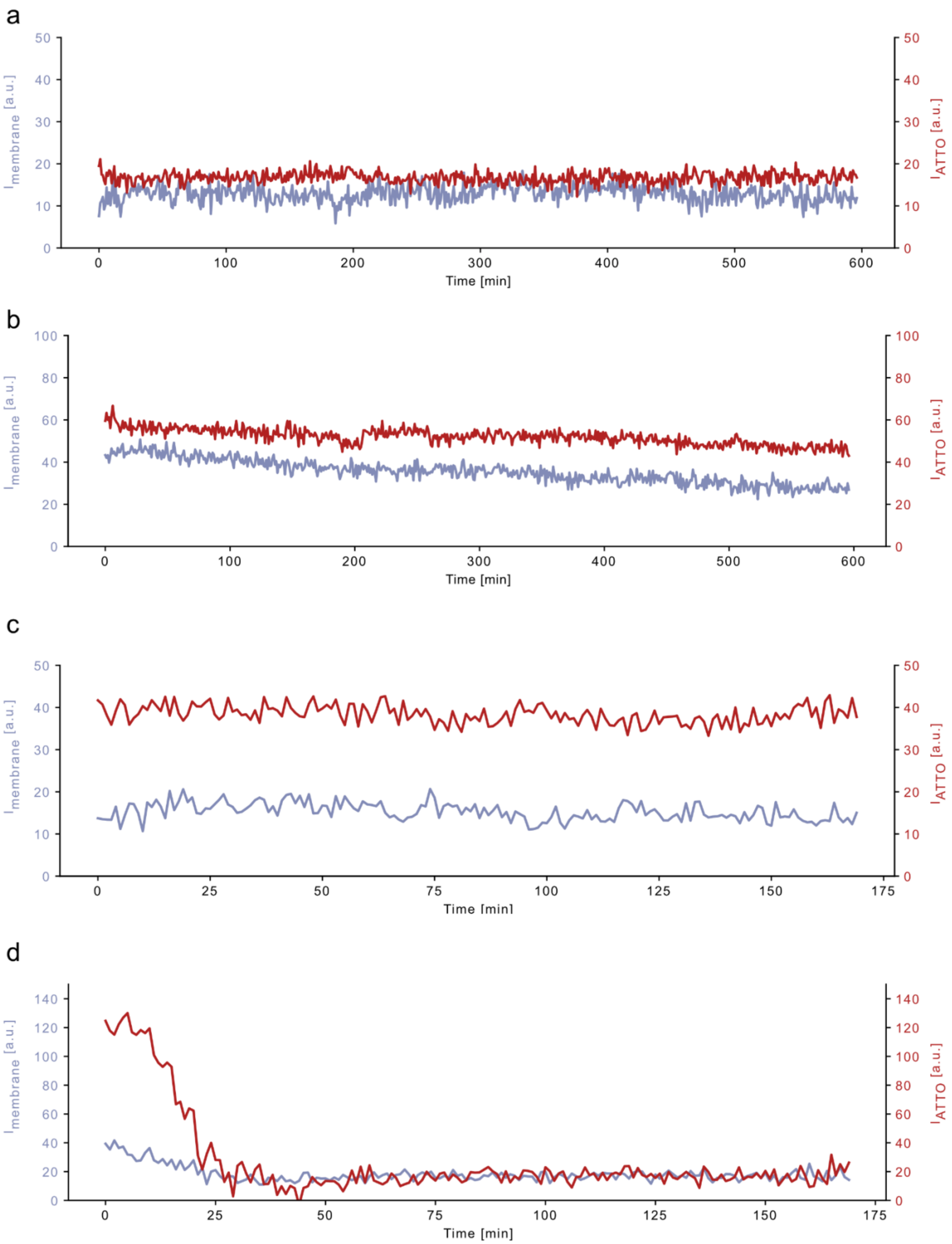
a
$I_{membrane}$ [a.u.]
$I_{ATTO}$ [a.u.]
Time [min]
b
$I_{membrane}$ [a.u.]
$I_{ATTO}$ [a.u.]
Time [min]
c
$I_{membrane}$ [a.u.]
$I_{ATTO}$ [a.u.]
Time [min]
d
$I_{membrane}$ [a.u.]
$I_{ATTO}$ [a.u.]
Time [min]

SI Fig. 12: Representative examples of real-time single-vesicle recordings with αSM demonstrating limited translocation. *99% DOPG liposomes with encapsulated ATTO655-carboxy, were monitored with a framerate of 1 image/min for 170 or 600 minutes. αSM were washed in the chamber after 7 minutes. Showing no membrane rupture (blue) for the two particles with a+b+c) no translocation events and d) multiple or constant leakage until the liposome was depleted. Both trajectories show a stable liposome membrane but do not allow for dyes' translocation through stepwise pore formation. The membrane signal and the ATTO655 signal are given in arbitrary units.*

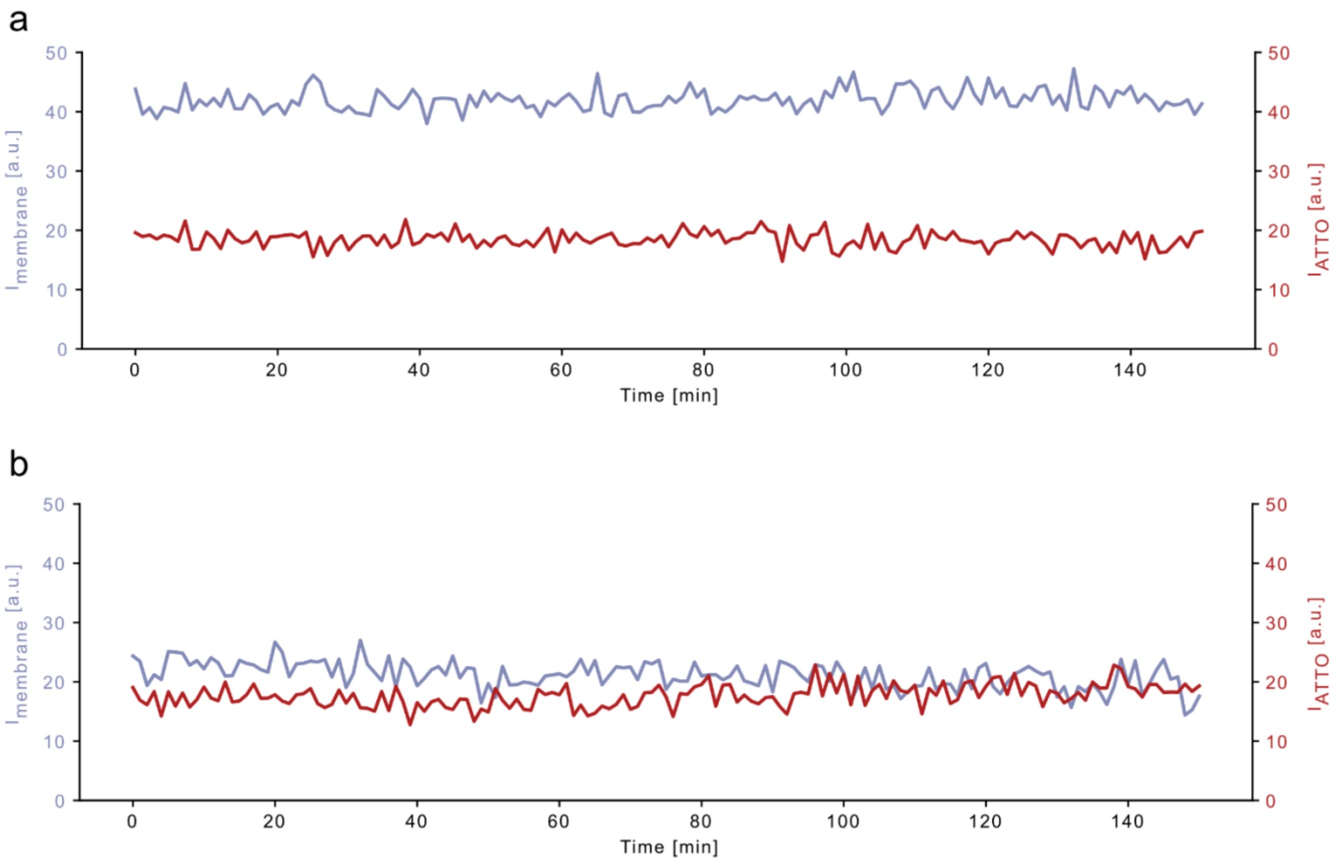


SI Fig. 13: Representative examples of real-time single-vesicle recordings with buffer only demonstrating no translocation. *99% DOPG liposomes with encapsulated ATTO655-carboxy, were monitored with a framerate of 1 image/min for 150 minutes. PBS buffer was washed in the chamber after 7 minutes. Showing no membrane rupture (blue) for the two particles with a+b) no translocation events. Both trajectories show a stable liposome membrane but do not allow for dyes' translocation. The membrane signal and the ATTO655 signal are given in arbitrary units.*

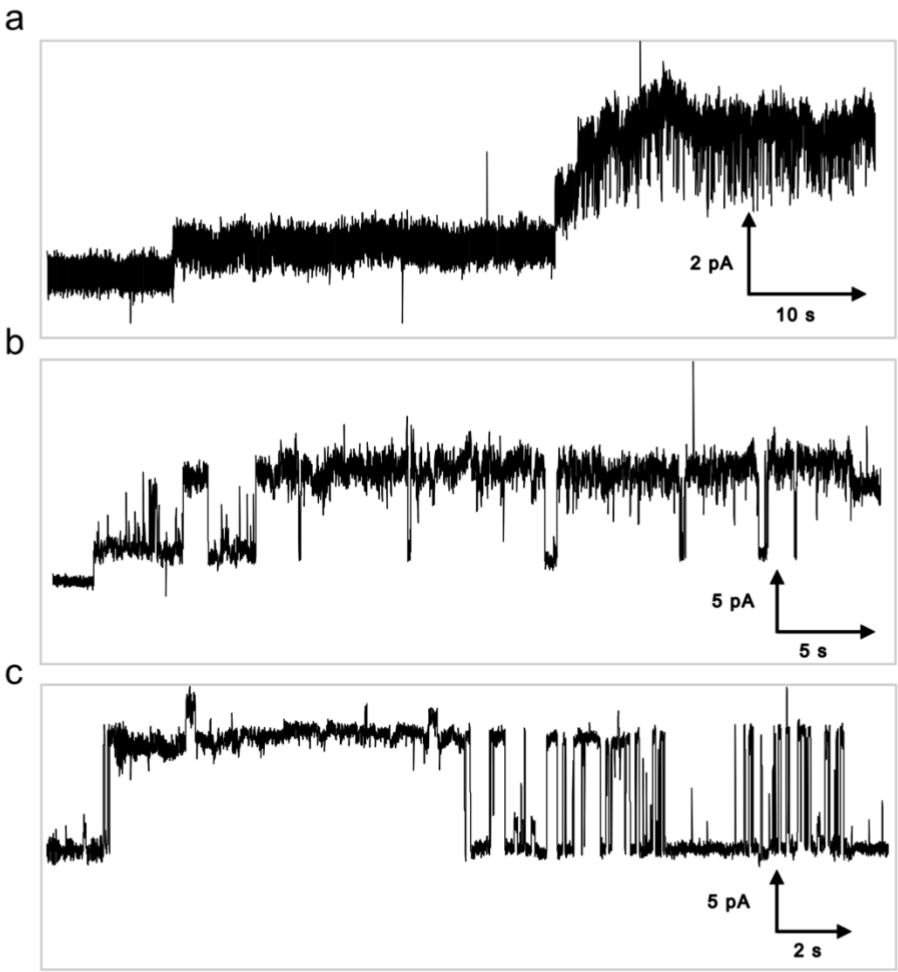


SI Fig. 14: Representative current trajectories showing the pore formation of α-syn oligomer on neutral-charged planar lipid bilayer. *These experiments were independently recorded at an applied voltage of 20 mV and in an electrolyte buffer: 1 M KCl, 50 mM Tris, pH 7.4. 1.4 ug of α-syn oligomer was added to both side of the chamber before recording. The neutrally charged planar lipid bilayer was formed by using 10 mg/mL DphPC.*

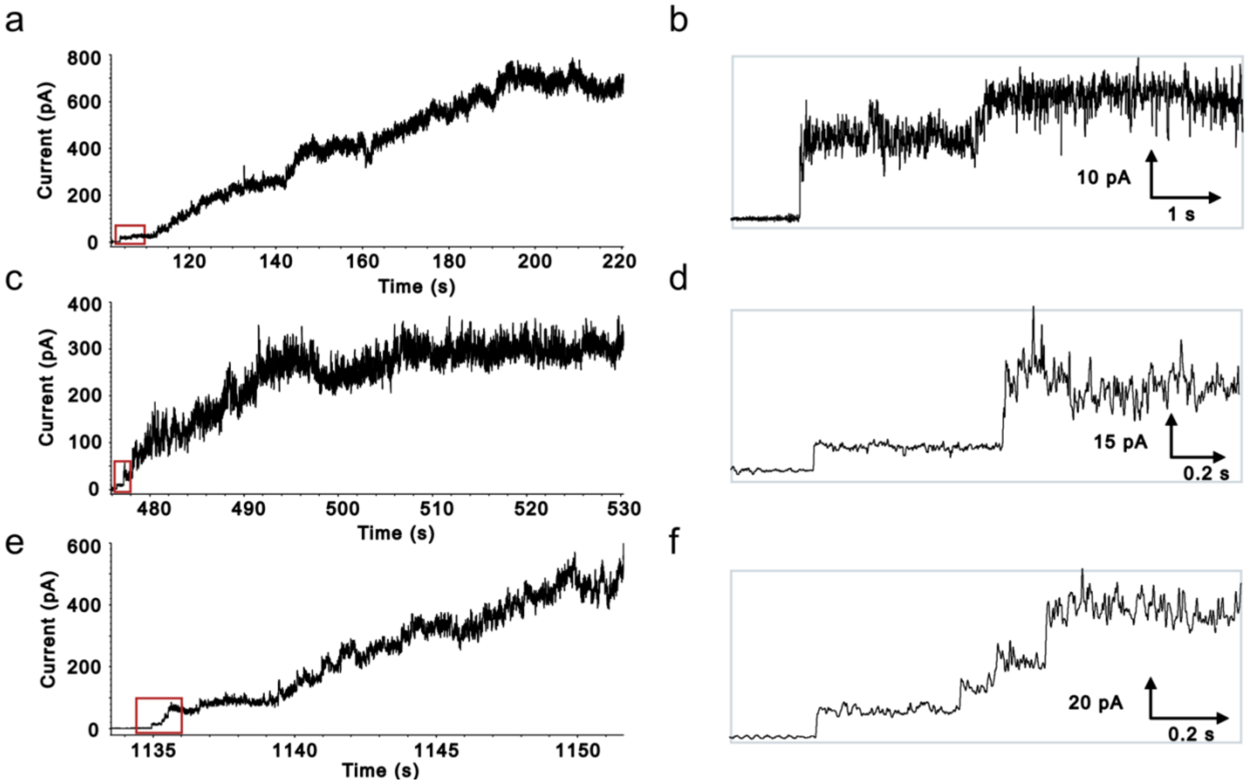


SI Fig. 15: Representative current trajectories showing the pore formation of α-syn oligomer on the negatively charged planar lipid bilayer. *These experiments were independently recorded at an applied voltage of 20 mV and in an electrolyte buffer: 1 M KCl, 50 mM Tris, pH 7.4. 1.4 ug of α-syn oligomer was added to each side of the chamber before recording. The negatively charged planar lipid bilayer was formed by using 10 mg/mL DphPG. a+c+e) trajectories of αSO insertion in negatively charged planar lipid bilayer. b+d+f) a zoom in of (a), (c) and (e).*

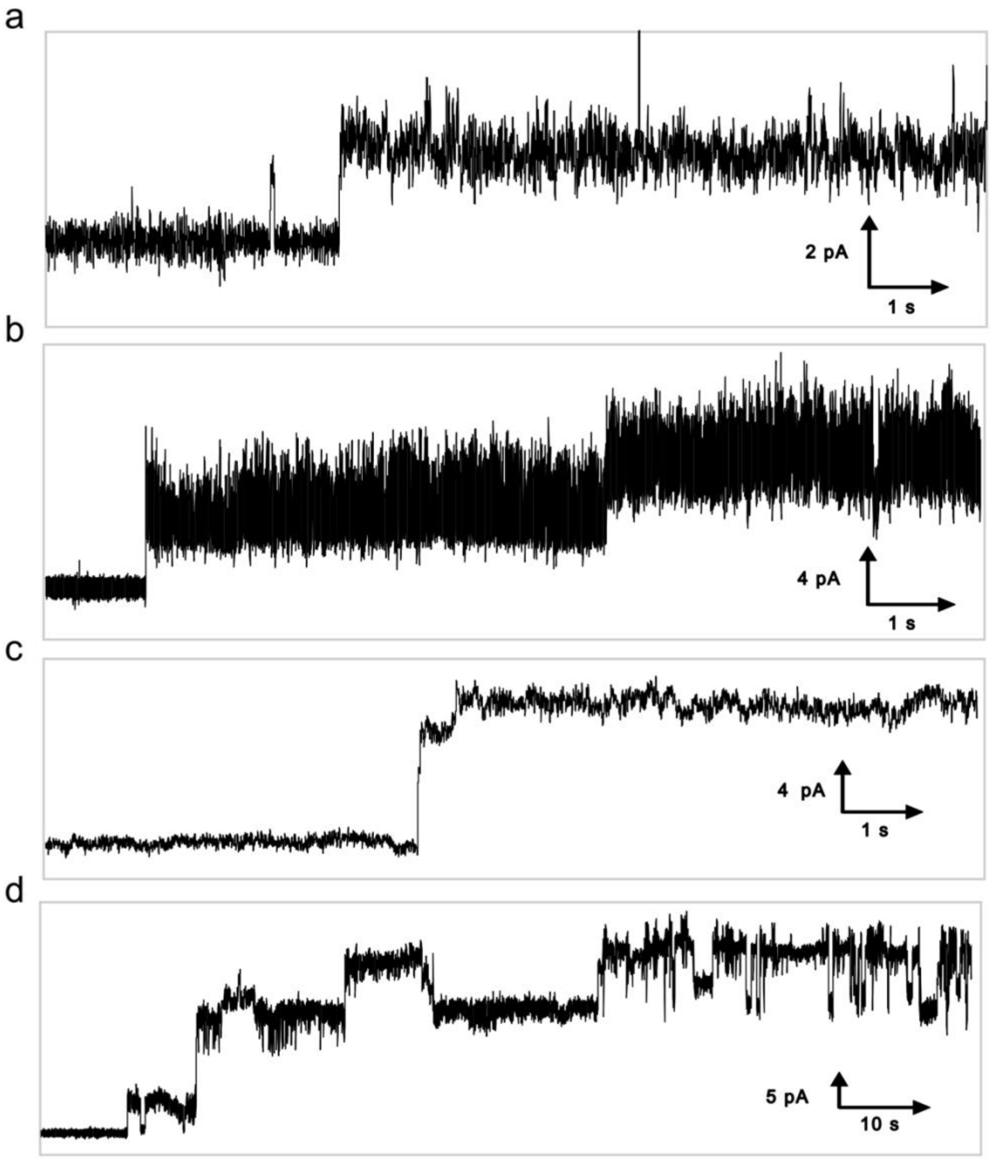


SI Fig. 5: Representative current trajectories showing the pore formation of α-syn oligomer on neutral-charged planar lipid bilayer in the presence of $CaCl_2$. *These experiments were independently recorded at an applied voltage of 20 mV and in an electrolyte buffer: 1 M KCl,1.5 mM $CaCl_2$, 50 mM Tris, pH 7.4. 1.4 ug of α-syn oligomer was added to each side of the chamber before recording. The neutrally charged planar lipid bilayer was formed by using 10 mg/mL DphPC.*

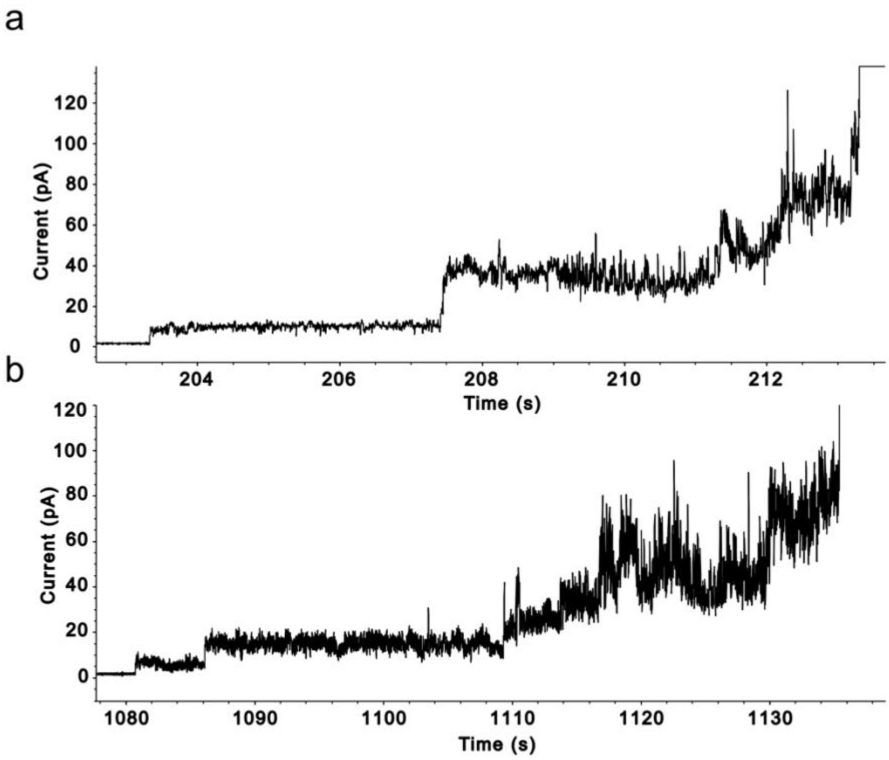


SI Fig. 17: Representative current trajectories showing the pore formation of α-syn oligomer on the negatively charged planar lipid bilayer in the presence of $CaCl_2$. *These experiments were independently recorded at an applied voltage of 20 mV and in an electrolyte buffer: 1 M KCl, 1.5 mM $CaCl_2$, 50 mM Tris, pH 7.4. 1.4 ug of α-syn oligomer was added to each side of the chamber before recording. The negatively charged planar lipid bilayer was formed by using 10 mg/mL DphPG.*

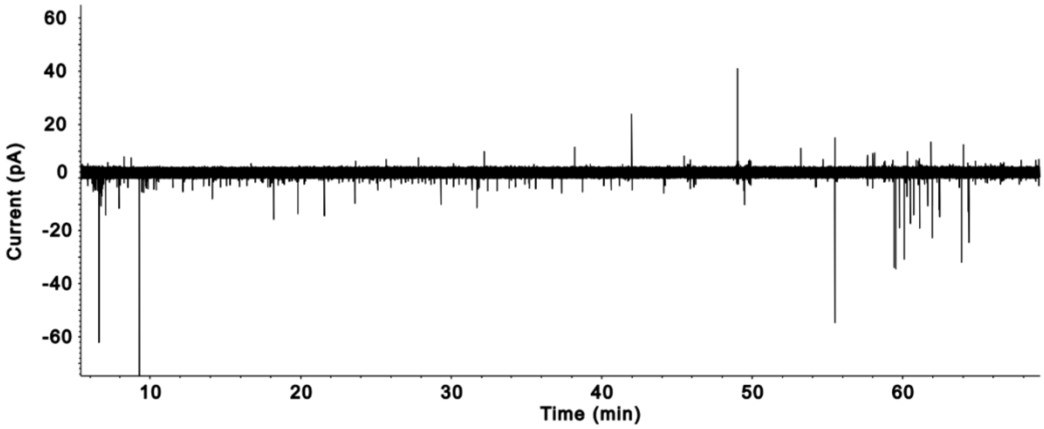


SI Fig. 18: Control experiment showing that α-syn monomer has no pore formation activity on the negatively charged planar lipid bilayer. *The experiment was recorded at an applied voltage of 20 mV and in an electrolyte buffer: 1 M KCl, 1.5 mM $CaCl_2$, 50 mM Tris, pH 7.4. 1.4 ug of α-syn monomer was added to each side of the chamber before recording. The negatively charged planar lipid bilayer was formed by using DphPG.*

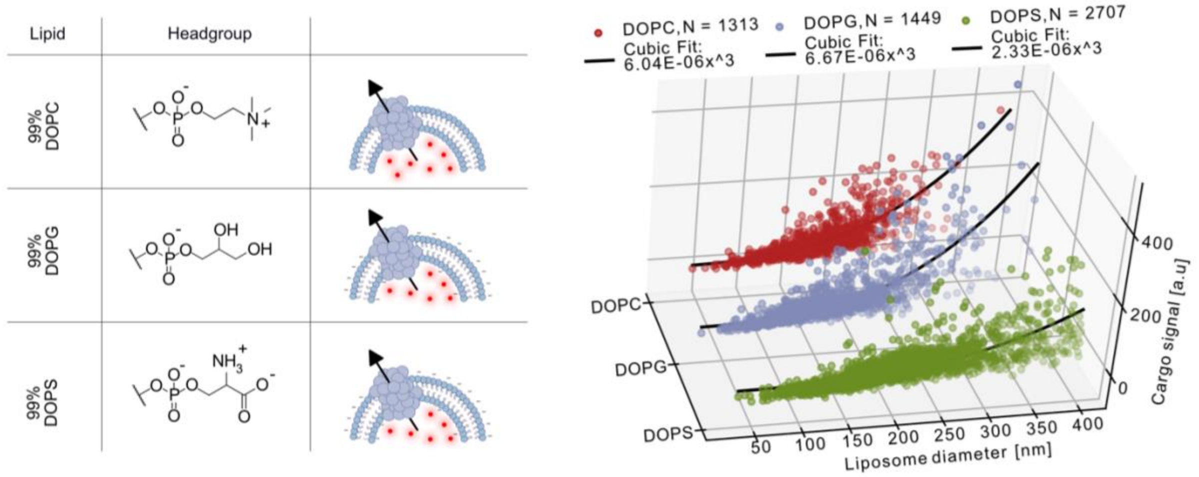


SI Fig. 19: Encapsulation of ATTO-655 carboxy in DOPC, DOPG and DOPS containing liposomes

*a) Liposomes are prepared with 3 different lipids. Negative lipids DOPS and DOPG and the neutral DOPC. b) Encapsulation efficiency of ATTO655-carboxy cargo dyes in the liposome lumen of DOPC, DOPS, and DOPG liposomes follows a cubic increasing encapsulation with increasing lumen size.*

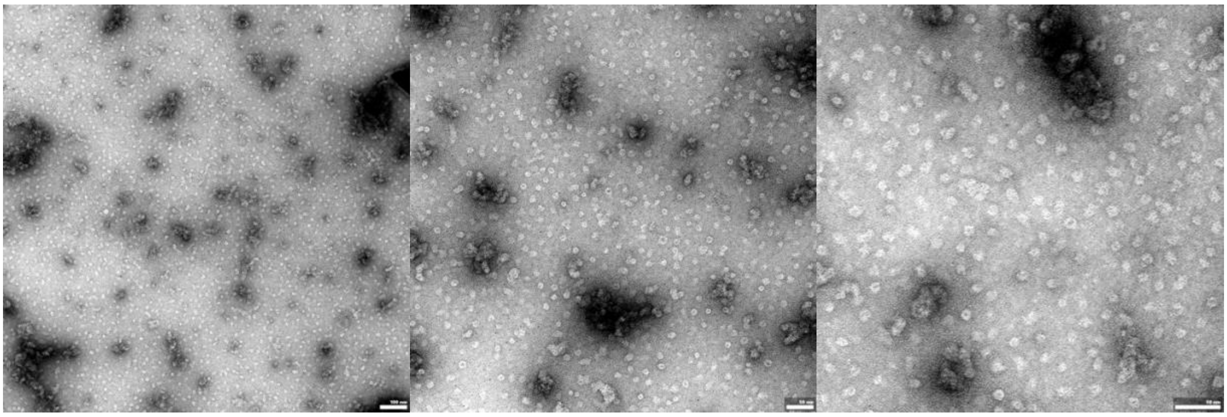

SI Fig 20. nsTEM images of α-syn oligomers

*Negative staining transmission electron microscopy (nsTEM) images of α-syn oligomers in absence of nanobodies. Scale bars are 100nm for the image to the right and 50nm for the middle and left image.*

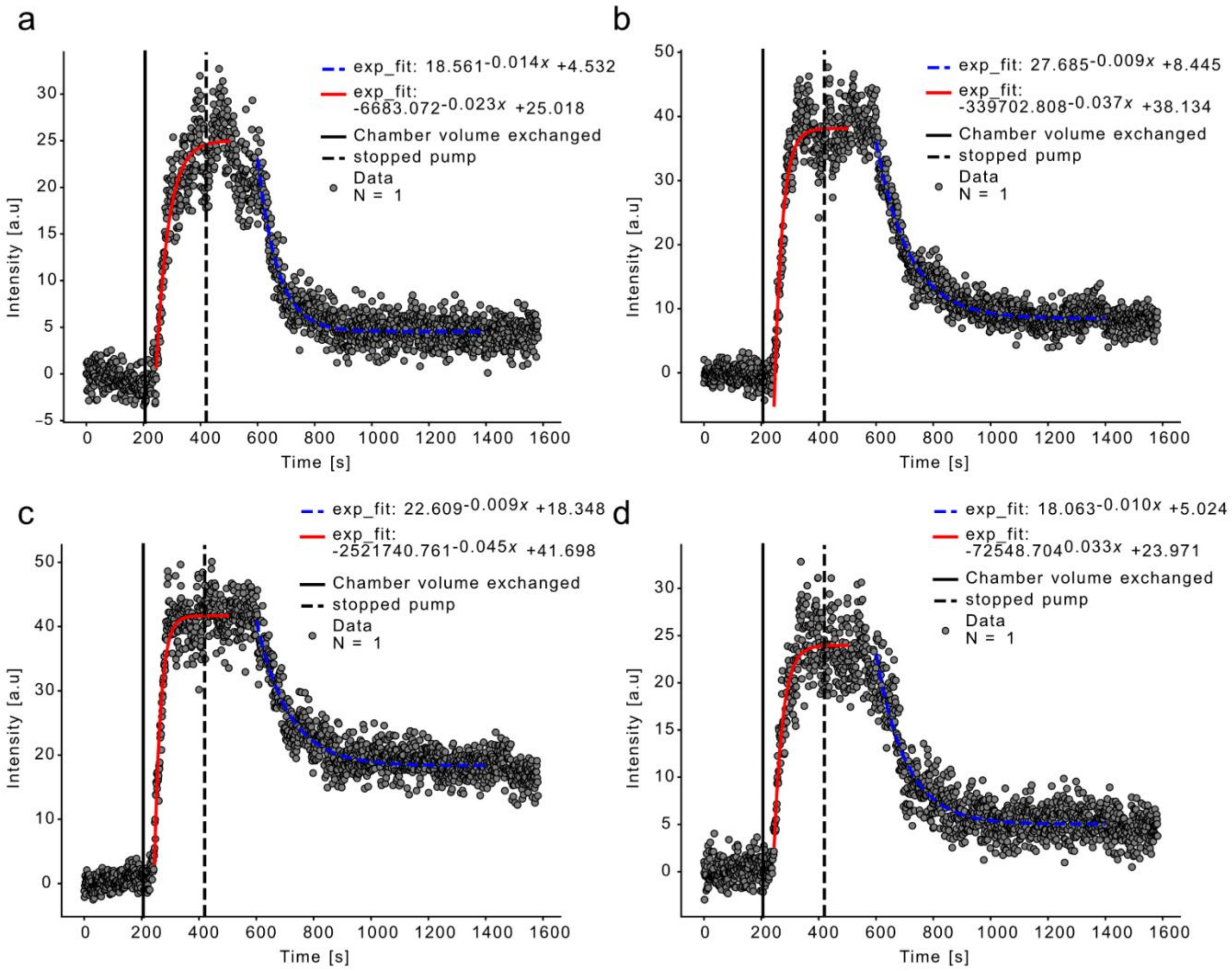


SI Fig. 21: Representative of real-time trajectories of αSO binding followed by NB1 binding to single-vesicles. *99% DOPG liposomes, were monitored with a framerate of 1 image/sec for 1600 seconds. After 180 seconds labeled αSO was pumped into the chamber and chamber volume was exchanged 20 seconds later (solid black line). The pumping stopped 4 minutes later (dotted black line), after membrane saturation. The recruitment of αSO (red slope) and NB1 is shown in liposomes with 4 different sizes: a) 30.76, b) 41.87, c) 69.63, and d) 54.86. Liposome size and αSO intensity are given in arbitrary units.*

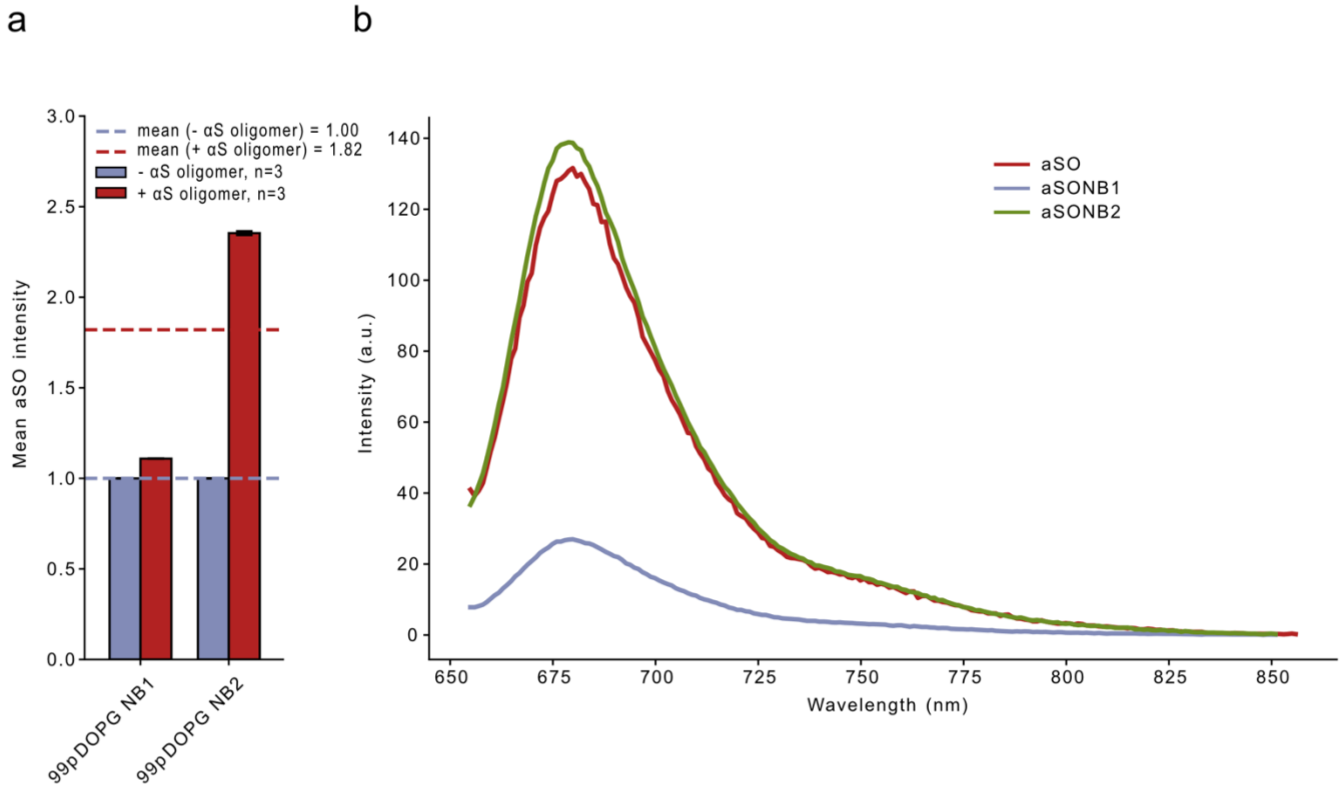


SI Fig. 6: Nanobody1 quenches ATTO655 signal. *a) Quantification of αSO recruited to the membrane shows a lower recruitment of oligomers to the membrane for oligomers bound to NB1. This observation was found to be due to signal quenching, and not lower recruitment. b) Fluorescent emission spectra of ATTO488-labled αSO (red), αSO:NB1 (blue), and αSO:NB2 (green). Spectra show that upon NB1 binding the fluorescent signal is quenched.*

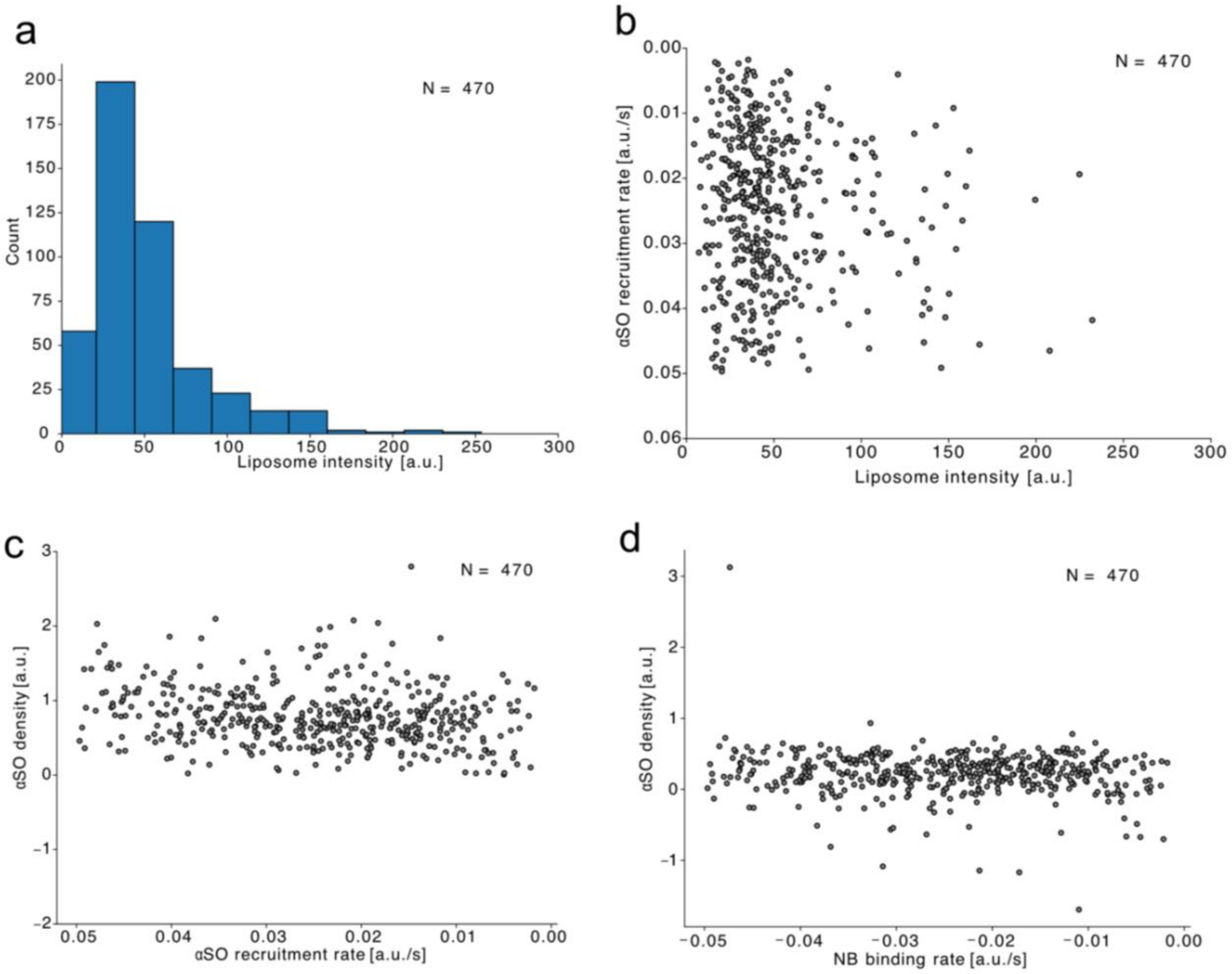


SI Fig. 23: Quantification of αSO recruitment and nanobody binding. *Fitted values from the equation in SI Fig. 20;* $\alpha SO_{intensity} = A * e^{B*x} + C$*, where A is the scaling factor, B is the αSO recruitment rate (or nanobody binding rate), and C is the αSO intensity at membrane saturation. a) Histogram of the liposome intensity (size [a.u.]) distribution shows the expected lognormal distribution. b) αSO recruitment rate from single liposomes plotted against the liposome size. c) αSO density (C / liposome size) plotted against the αSO association rate. d) αSO density plotted against nanobody binding rate.*

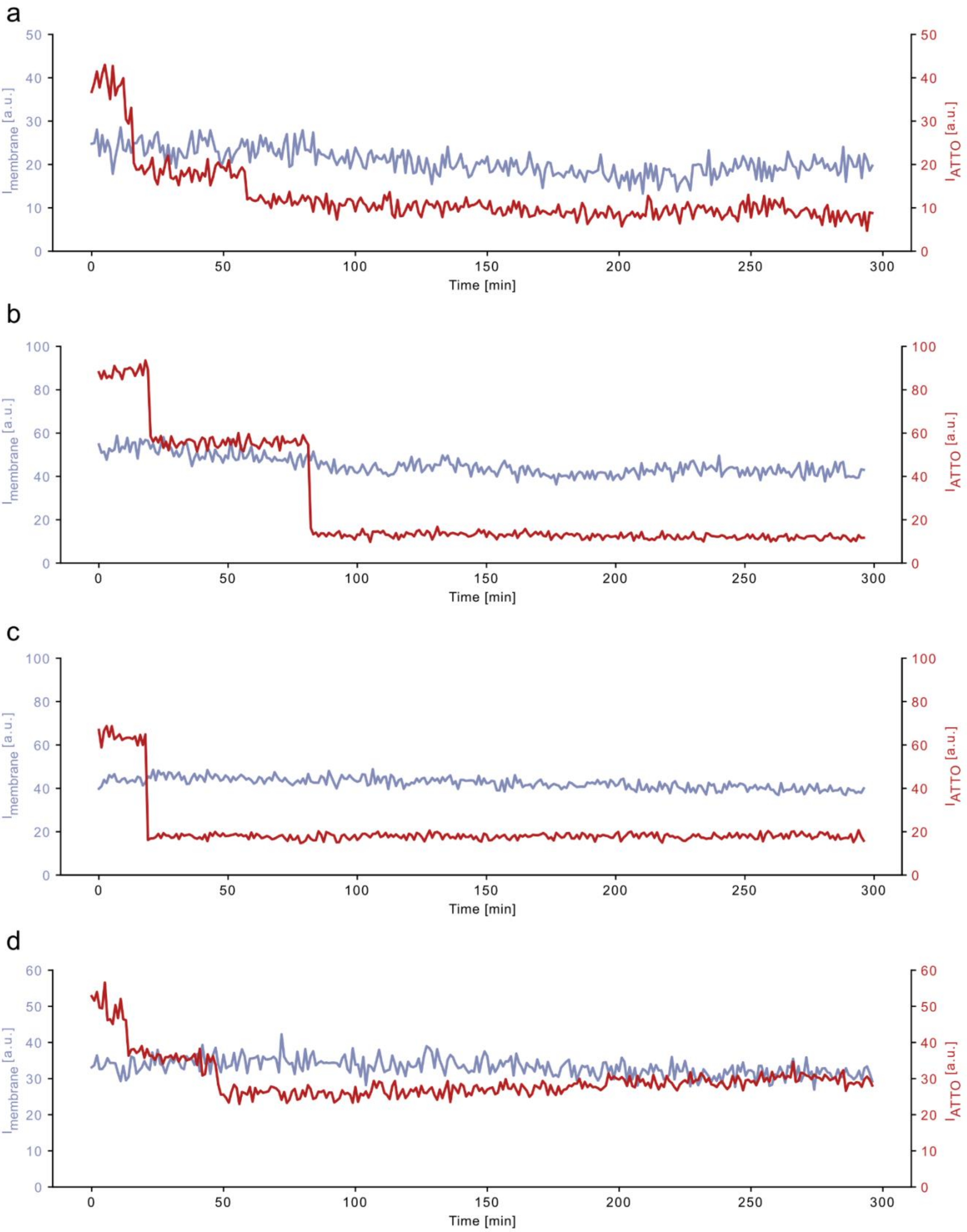


SI Fig. 24: Representative examples of real-time single-vesicle recordings demonstrating αSO pore formation in presence of NBs. 99% DOPG liposomes with encapsulated *ATTO655-carboxy*, were monitored with a framerate of 1 image/min for 300

minutes. a+b) αSO:NB1 and or c+d) αSO:NB2 were washed in the chamber after 7 minutes. Showing no membrane rupture (blue) for the two particles with a+b+d) two translocation events and c) one translocation event. All trajectories show a stable liposome membrane and a dynamic pore formation with two distinct open pore formations allowing for dyes' translocation. The membrane signal and the *ATTO655 signal* are given in arbitrary units.
